# Planetary-scale metagenomic search reveals new patterns of CRISPR targeting

**DOI:** 10.1101/2025.06.12.659409

**Authors:** Simon Roux, Uri Neri, Brian Bushnell, Brayon Fremin, Nikhil A. George, Uri Gophna, Laura A. Hug, Antonio Pedro Camargo, Dongying Wu, Natalia Ivanova, Nikos Kyrpides, Emiley Eloe-Fadrosh

## Abstract

Interactions between microbes and their mobile genetic elements (MGEs), including viruses and plasmids, are critical drivers of microbiome structures and processes. CRISPR-Cas systems are important regulators of these host-MGE interactions, but a global understanding of CRISPR loci diversity, activity, and roles across Earth’s biomes is still lacking. Here, we use an optimized computational approach to search short-read data and collect ∼800 million CRISPR spacers across ∼450,000 public metagenomes. From this extensive CRISPR spacer dataset, we identified 1.18 billion hits between 41 million spacers and 2.5 million viruses and plasmids. Focusing on the role of CRISPR as anti-phage defense, we observed an over-representation of prevalent and conserved spacers consistent with a positive selection pressure associated with phage targeting. Identification of sustained CRISPR targeting by microbes not expected to be viable hosts suggested that some phage genomes may frequently enter non-host microbial cells. Taken together, these results outline the extensive diversity of CRISPR loci across microbiomes, and highlight key parameters driving CRISPR targeting patterns likely to influence strain-level diversification and selection processes within microbial populations.

## Introduction

Microbes are the dominant form of life on Earth, driving major biogeochemical cycles and shaping the biology of larger plants and animals, including humans^1,2^. Microbes most often coexist in complex communities, known as microbiomes, in which interactions between organisms are critical drivers of microbial ecology, evolution, and metabolic activity^3,4^. Among these, interactions between microbes and mobile genetic elements (MGEs), including plasmids and viruses, can shape long-term genetic exchange and evolutionary trajectories of microbial populations. Viruses in particular are now broadly recognized as key drivers of microbial population structure, diversity, and metabolism^5–8^. Yet the full breadth of interactions between microbes and MGEs are still only partially explored, and eco-evolutionary drivers of MGE-host dynamics remain to be characterized.

CRISPR-Cas systems, encoded in ∼40% of bacterial and ∼85% of archaeal genomes^9^, play a key part in these MGE-host interactions. Since their original discovery, CRISPR-associated enzymes have quickly become a foundational part of the genome editing toolkit as they can recognize and cleave specific DNA or RNA templates ^10–12^. In nature, CRISPR-Cas systems typically target exogenous DNA or RNA molecules, and store short fragments derived from these templates, termed “spacers”, to enable recognition and targeting of similar exogenous elements in future encounters^9,13–16^. As CRISPR spacers are stored in dedicated and identifiable loci (“arrays”) in microbial genomes, these can be used to survey past encounters between microbial populations and other organisms or genetic elements, especially plasmids and viruses. So far, the diversity of spacer arrays and the impact of spacer targeting on MGE populations have been mostly studied *in vitro* or in individual habitats^17–21^. Nonetheless, the first studies of CRISPR spacers obtained from shotgun metagenomes, i.e. direct sequencing of DNA extracted from a complex microbiome, suggested that CRISPR arrays in natural communities are broadly but unevenly distributed across taxa and environments, that some spacer arrays are highly diverse across space and time, and that spacers can target both sympatric and allopatric viruses^20,22–26^.

As these metagenomic studies have been performed on relatively small-scale datasets, several open questions remain regarding the overall diversity and MGE targeting of CRISPR arrays^21,27–31^. First, several tools have been designed to extract CRISPR spacers from metagenomes, which sometimes led to conflicting observations, ranging from very high population diversity and spacer turnover to highly stable arrays with low rates of new spacer acquisition, without a clear indication of whether these reflect biological or methodological differences^21,30,36^. Second, several metagenomic studies recently reported viruses targeted by bacteria assigned to different classes or phyla, far beyond the host range of characterized virus isolates^37–39^. This suggests that either some uncultivated viruses are able to infect hosts across these large phylogenetic distances, or CRISPR targeting may occur between viruses and non-host microbial populations^38^.

Here, we present an approach to systematically extract CRISPR spacers from short reads across 467,018 public metagenomes and build a global database of 794,125,610 CRISPR spacers connected to 274,000 distinct repeats. We leverage this unique dataset to explore global diversity and activity patterns of CRISPR systems across ecosystems, identify different levels of CRISPR targeting presumably associated with different types of microbe-MGE interactions, and use this new framework to re-interpret previous observations of viruses targeted by CRISPR arrays from phylogenetically distant microorganisms.

## Results

### Metagenome reads encode an extensive diversity of CRISPR spacers

To evaluate the global diversity of CRISPR spacers across taxa and environments, we first used CRISPRCasTyper^40^ to predict CRISPR arrays across 383,318 public genomes and 33,727 metagenomes. After quality control and curation (see Methods and Supplementary Data File 1), a total of 581,889 non-redundant CRISPR repeats were retained (Fig. 1A). Most of these predicted repeats were only detected on short metagenome contig with limited (if any) genomic context (Fig. 1B). Among repeats that could be classified taxonomically (n=109,810), most (61.3%) were assigned to a single family or genus, spanning across 159 phyla, 2,481 families, and 8,418 genera of bacteria and archaea (Fig. 1B, Fig. S1).

**Fig. 1.**
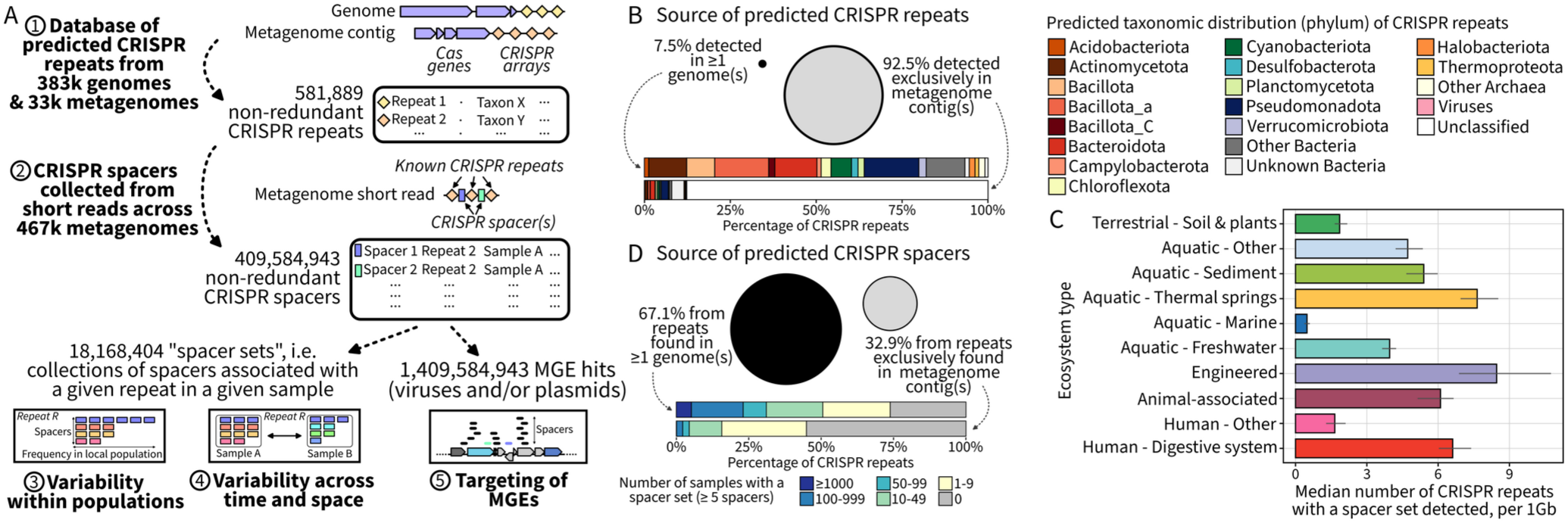
Large-scale detection of CRISPR spacers across metagenomes. **A.** Schematic of the approach used to collect and analyze a global CRISPR spacer dataset. Potential CRISPR repeats were first identified from assembled genomes and metagenomes, and then used in the SpacerExtractor tool to collect spacers from metagenomic reads. Spacers were then analyzed in terms of diversity within and between samples, and matched to viruses and plasmids (IMG/VR and IMG/PR) to detect potential targeting. **B.** Distribution of the predicted CRISPR repeats source (metagenome only, or detected in ≥1 genome(s)), and taxonomy. The relative proportion of each CRISPR repeats source is indicated by the size of the circle, and the taxonomic distribution for each source shown in a bar chart underneath for each group. **C.** Median number of distinct repeats for which spacers were identified for individual metagenomes, per 1Gb of metagenomic reads, across ecosystems. The colored bar represents the average value obtained across 50 random subsamples of 1,000 metagenomes for each category, and the error bar spans from the minimum value to the maximum value observed across these subsamples.

We next leveraged this extensive database of predicted CRISPR repeats with a new pipeline designed to efficiently extract CRISPR spacers from short reads (“SpacerExtractor”, see Methods, Supplementary Text 1, Fig. S2) across 467,018 metagenomes publicly available in NCBI’s SRA (Fig. 1A, Supplementary Data File 2) ^41^. After quality control and denoising (see Methods and Fig. S3), this search yielded a total of 794,125,610 spacers, which dereplicated into 409,584,943 distinct spacers associated with 274,000 repeats across 367,609 metagenomes. After controlling for sequencing depth, a higher number of CRISPR repeats was detected on average in wastewater, bioreactor, and hot spring microbiomes, while a lower number was observed in marine samples, likely reflecting a combination of differences in community complexity and frequency of CRISPR array in these populations^23^ (Fig. 1C, Fig. S4). The taxonomic distribution of these CRISPR repeats was consistent with expected patterns of bacterial taxa distribution, with e.g. *Lachnospira* and *Bifidobacterium*-assigned repeats being among the most frequently detected in human gut samples, and *Microcystis* the most common in freshwater (Fig. S5). Overall, spacers were typically (99.0%) associated with a single repeat sequence, or associated with a single genus when linked to multiple repeat sequences (78.5%).

Spacer sets, defined here as sets of 5 or more non-redundant spacers associated with a given repeat in a given sample (Fig. 1A), could be readily obtained for most repeats identified in at least one genome ≥ 10 samples (Fig. 1D). Conversely, spacer sets for repeats exclusively detected on metagenomic contigs were typically only found in <10 samples, and nearly half (55.1%) of these repeats were not associated with any spacer set (Fig. 1D). These metagenome-only repeats are thus likely a combination of mispredictions, i.e. repetitive sequences not part of a genuine CRISPR-Cas system, and CRISPR-Cas systems found on rare and under-sampled genomes. Ultimately, despite representing only 7.5% of the repeat database, predicted CRISPR repeats found in at least one genome yielded 67.1% of all spacers, meaning that most of the spacer dataset is associated with known CRISPR-Cas systems in bacteria and archaea (Fig. 1D, Fig. S1). Taken together, these results illustrate that the combination of an extensive CRISPR repeat database and large-scale metagenome mining provides a uniquely broad picture of the global diversity of CRISPR spacers across microbiomes.

### CRISPR spacers show high turnover rate and limited persistence across samples

We next explored the content and diversity of 18.1 million spacer sets (5 or more spacers), corresponding to spacers associated with 274,000 unique repeats across 367,609 metagenomes (Fig. 1A). Specifically, we compared the composition of these spacer sets within and between samples to evaluate the prevalence and persistence of individual spacers in natural microbial populations (Fig. S6).

Overall, most (52.5%) spacer sets included ≥20 distinct spacers, with 1,346,554 (7.4%) including ≥100 distinct spacers (Fig. 2A). Among sets with sufficient sampling (≥1 spacer(s) with ≥20x coverage, n=2,068,538 sets), 75.7% showed a minority of high-abundance or “common” spacers, presumably shared across most members of the population, followed by a long tail of rare spacers (Fig. 2B). Accordingly, across the entire dataset, a two-third (67.0%) majority of spacers were singletons, i.e. identified on a single read (Fig. S7). This is consistent with a previous survey that used targeted PCR to deeply sequence the spacer diversity associated with a known repeat, and observed a large majority of low-frequency spacers^36^. Clustering of spacers at lower identity levels suggested that more than half (62.9%) of the singletons were not closely related to another spacer in the same set, and thus are likely to represent genuinely rare spacers as opposed to sequencing artifacts (Fig. 2C, Supplementary Text 2). Taken together, these results suggest that, at any given time, individual cells within a microbial population encode a small subset of spacers shared broadly within the population, next to a larger number of rare spacers found in a minority of population members. Across our dataset, the largest spacer sets were observed in thermal springs and engineered ecosystems, and for uncultivated taxa known or predicted to engage in microbe-microbe interactions, possibly reflecting specific eco-evolutionary pressures leading to high population diversity at CRISPR loci (Supplementary Text 2, Fig. S8).

**Fig. 2.**
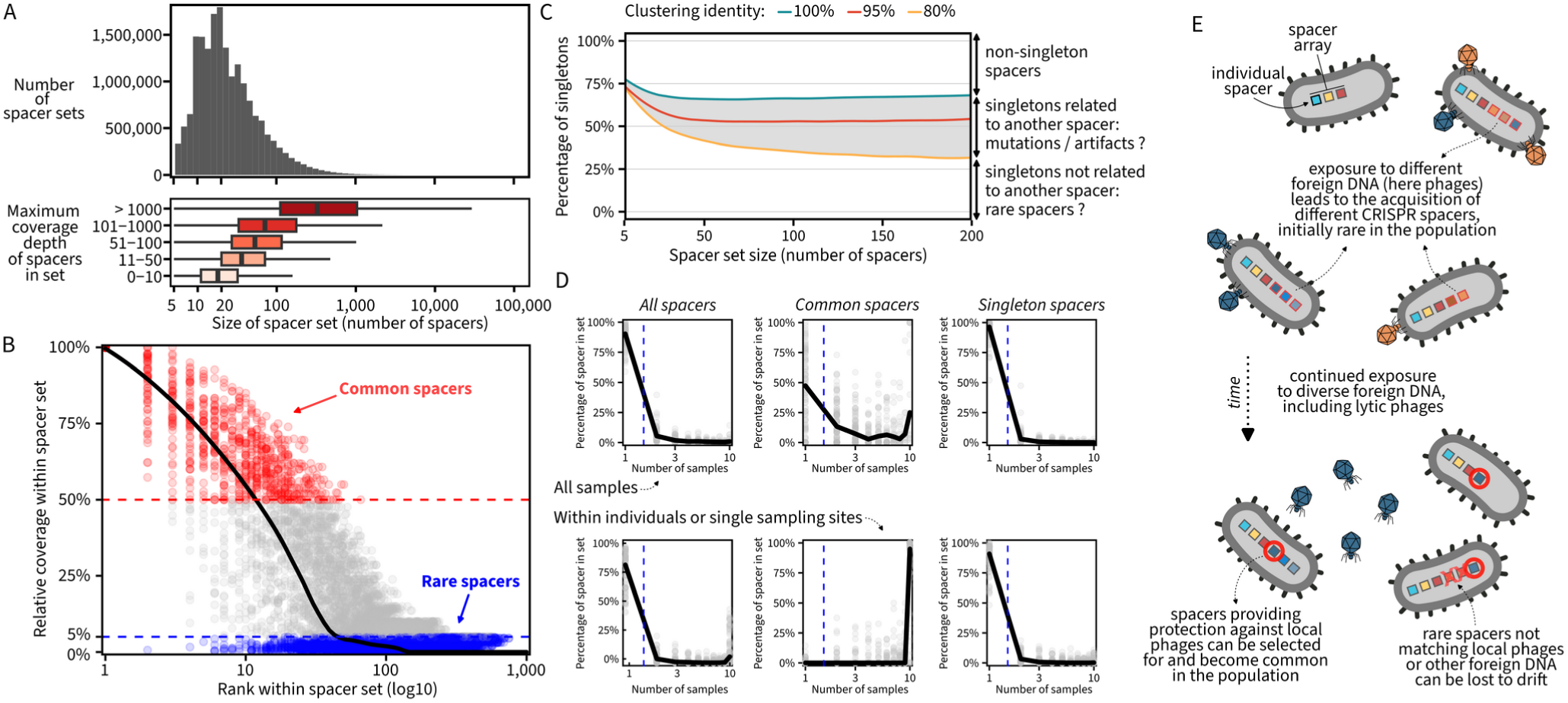
Spacer sets diversity within and between samples. **A.** Distribution of the size of spacer sets, i.e. of the number of non-redundant spacers associated with a given repeat in a given sample. The bottom panel shows the distribution of spacer set size as boxplots for different set coverages, estimated here based on the maximum spacer coverage in the set. The box shows the inter-quartile range (IQR) with the median highlighted by a black line, and whiskers extend to 1.5*IQR. **B.** Illustration of the typical distribution of spacer coverage within collected spacer sets. The plot is based on a subsample of 100 spacer sets per ecosystem, each with at least 1,000 spacer-containing reads and at least one spacer with ≥20x coverage. Each spacer coverage is then plotted according to its rank (x-axis, log10) and its coverage relative to the maximum coverage in the set (y-axis), with common (≥50% of the maximum coverage) and rare (<5% of the maximum coverage) spacers highlighted in color. The black curve is based on a local polynomial regression across all data points. **C.** Percentage of singleton spacers (i.e. spacers identified on a single read, y-axis) within set as a function of the set size (x-axis) and minimum identify percentage used for clustering (in colors). The plot shows the median percentage of singletons for each spacer set size, for sets from 5 to 200 spacers. The distribution of singleton percentage for categories of spacer set size are provided in Fig. S7. **D.** Number of samples in which individual spacers were detected for individual repeats, for all spacers (left), common spacers only (middle) or singleton spacers only (right). The number of samples is log10-transformed (x-axis), and a dashed blue line indicates the border between one sample (left) and multiple samples (right). Plots are based on a random subset of 10 samples for repeats found in 10 samples or more. The top row shows plots based on any random subset of 10 samples, while the bottom row shows the same plot for subsets of 10 samples associated with the same subject (for host-associated samples) or location (for environmental samples). **E.** Schematic overview of the potential mechanisms that could lead to a high observed diversity for CRISPR loci both within a sample and between samples.

CRISPR spacers are also expected to vary across time and space as new spacers are acquired and potentially selected for. To evaluate spacer variability across samples, we focused on the 17,585 repeat sequences with high-coverage sets (≥1 spacer(s) with ≥20x coverage) available for 10 or more samples, and compared the composition of these sets across samples (Fig. S6). Most spacers (86.5%) were only detected in a single sample, although spacers identified as common in the population for at least one sample were more frequently identified in multiple samples (∼53.7%, Fig. 2D). When considering only samples from a single subject (for host-associated ecosystems) or location (for environmental samples), a similar percentage (84.7%) of spacers were detected in a single sample, but almost all common spacers (95.6%) were detected in multiple samples (Fig. 2D). This is consistent with recent analyses of CRISPR loci diversity in individual human gut, shale wells, and marine aggregate microbiomes^21,27,29,30,42^, and suggests that (i) new spacers are frequently acquired by individual cells, leading to high alpha and beta diversity at CRISPR loci, (ii) most newly-acquired spacers are quickly lost to drift, and only a minority is retained and becomes common in the population, likely because of direct selection or genetic draft, and (iii) these common spacers are especially conserved through time within stable and established microbiomes (sensu ^43^) such as in a single subject or location (Fig. 2E). Opposing this overall trend, a small number of repeats (∼2.0%) were associated with an unusually low diversity of spacers within and between samples, and may represent “memory arrays” as previously suggested ^44^ (Fig. S9, Supplementary Text 2).

### CRISPR spacers frequently match multiple potential targets

Given this expanded set of CRISPR spacers, we next looked for potential protospacers, i.e. spacer hits, in two large databases of viruses (IMG/VR) and plasmids (IMG/PR). Considering only hits covering the entire spacer with 0 or 1 mismatch, we identified spacer hits for 36.1% of IMG/VR sequences (n=2,011,412), including 81.8% of high-quality tailed phages genomes (HQ *Caudoviricetes*, n=290,883), for a total of 1.18 billion hits across 38 million distinct spacers (Fig. 3A, Fig. S10). Beyond *Caudoviricetes*, 9 million hits were observed to other phages including 4 million hits to *Malgrandviricetes* ssDNA phages, providing new information on the potential host range of these and other atypical viruses (Supplementary text 3, Fig. S11). Similarly, a majority (59.9%) of plasmids in IMG/PR had ≥1 hit(s), including 84.8% of the predicted complete plasmids and 79.5% of plasmids encoding a mating pair formation system, for a total of 212 million hits across 3 million spacers (Fig. 3A, Fig. S10). These spacer hits were more frequently observed between spacers and viruses or plasmids from similar ecosystems (Fig. S10), and represent a 6-fold and 2-fold increase compared to the current data in IMG/VR and IMG/PR, respectively. Based on these hits, potential Protospacer-Adjacent Motifs (PAMs) could be predicted for 68.2% of repeats based on sequence conservation around the spacer hits (Fig. 3B). When predicted, the PAM sequences were overall consistent with known motifs described in the literature for the corresponding CRISPR types^45,46^ (Fig. 3B, Fig. S12, Table S1).

**Fig. 3.**
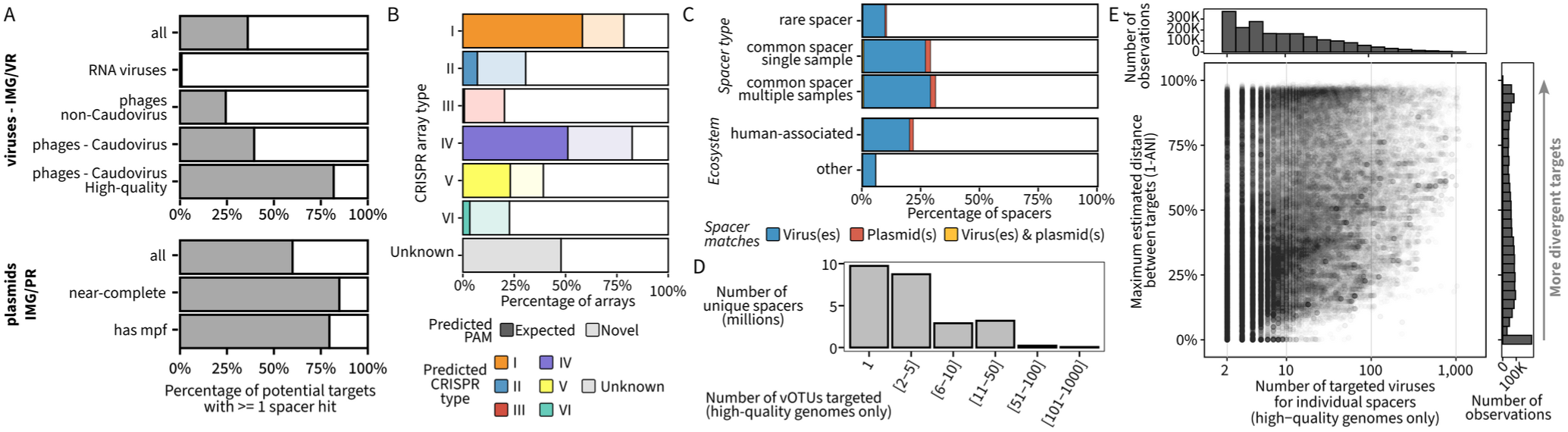
Spacer hits in IMG/VR (viruses) and IMG/PR (plasmid) databases. **A.** Percentage of virus genomes (top) and plasmids (bottom) with at least 1 spacer hit (100% spacer coverage, 0 or 1 mismatch). For both viruses and plasmids, the percentage across the entire IMG/VR or IMG/PR database is first indicated, followed by percentages for specific subsets. Caudovirus are sequences assigned to the class *Caudoviricetes*. “High-quality” is used to designate genomes predicted to be ≥90% complete. “mpf” refers to mating pair formation genes. **B.** Prediction of protospacer-adjacent motifs (PAMs) based on sequence conservation around spacer hits to IMG/VR and IMG/PR, by predicted CRISPR array type. Predicted array types were obtained from CRISPRCasTyper. For each array type, the percentage of arrays (i.e. unique repeats) for which a putative PAM was identified is in color, with solid colors when this predicted PAMs matched a known motif previously described in the literature, and light color otherwise. A more detailed description of the predicted PAMs is available in Fig. S12. **C.** Percentage of unique spacers matching a sequence in IMG/VR (viruses), IMG/PR (plasmids), or both. This percentage is shown for different categories of spacers based on their prevalence within and between samples (top), or their ecosystem of origin (bottom). The percentage of hits for common spacers was significantly greater than the one for rare spacers, and the percentage of hits for spacers from human-associated samples was significantly greater than the one for spacers from other ecosystems (2-sample test for equality of proportions, p-value <2.2e-16). **D.** Distribution of the number of vOTUs matched by individual spacers. For this analysis, only matches to high-quality virus genomes were considered, as including genome fragments may artificially inflate the number of vOTUs due to incomplete clustering. **E.** Genome diversity of targets for spacers matching 2 or more viruses. A subsample of 2,000,000 spacers matching ≥2 high-quality virus genomes was selected, and the similarity between target genomes evaluated for each spacer. The y-axis represents the maximum distance (i.e. 1 – average nucleotide identity, ANI) observed between targets for a given spacer. The distribution of the number of observation along the x-axis and y-axis are based on the 2,000,000 spacers, and a random subset of 100,000 spacers are included in the x-y plot for clarity.

Matches to viruses and plasmids were more frequently observed for common spacers (Fig. 3C, 30.5% with hit(s) compared to 10.1% for rare spacers), and spacers from human-associated samples (21.8% vs 5.64% on average for other ecosystems), the latter likely reflecting a database bias towards human microbiome spacers, viruses, and plasmids (Fig. 3C). Strikingly, among spacers with at least one hit, 60.8% matched two or more distinct virus genomes, designated in IMG/VR as virus operational taxonomic units or vOTUs (Fig. 3D). These identical matches across distinct vOTUs were found across both closely related and mostly unrelated viruses (Fig. 3E), and more frequently associated with genes with known functions (47.9% of hits compared to 40.9% for spacers matching a single vOTU, Table S2). This is consistent with previous observations from smaller-scale studies of spacers matching conserved genes and genes laterally exchanged between viruses^29,30^, and could reflect arms race dynamics in which spacers targeting the most critical and common regions of virus genomes would be preferentially selected for and retained in CRISPR arrays. However, the presence of these spacers matching multiple and sometimes unrelated viruses also means that, in order to reliably understand ongoing virus-host interactions, individual spacer hits may be misleading and a combined analysis of all spacer hits from a given repeat to a given target is required.

### Diversity-generating retroelements (DGRs) are associated with sustained targeting by non-host microbes

Given this high frequency of spacers potentially matching multiple targets, we next explored whether analyzing all spacers connecting a given repeat to a given virus could be used to distinguish virus-host interactions from spurious and non-specific hits. To that end, we focused on the 314,567 near-complete and complete viral genomes (“HQ” viruses) with at least one spacer hit, and leveraged 184,808 viruses with known hosts, either obtained from isolates or detected as a provirus (Supplementary Text 4).

We first focused on the number of distinct spacer hits identified between a given repeat and a given virus. Across 16,424,746 pairs in total, 73.7% included only a few spacer hits (<10 spacers), while 26.3% included ≥10 spacer hits (0 or 1 mismatch between spacer and virus, Fig. 4A, Fig. S13). For viruses with known hosts, the former cases were frequently (54.1%) associated with non-host taxa and may derive in part from spurious spacer alignments^47^, while targeting and host taxa were overwhelmingly consistent for cases with ≥10 spacer hits (76.8%, Fig. 4B). This indicates that the number of distinct near-exact (0 or 1 mismatch) spacer hits, possibly combined with the total number of base pairs covered by spacers on the virus (Fig. S14), are useful metrics to identify CRISPR targeting associated with true virus-host pairs. Across all HQ viruses, nearly half (48.1%) showed at least one repeat with ≥10 near-exact spacer hits, and may thus be connected to a putative host even when using this relatively conservative cutoff.

**Fig. 4.**
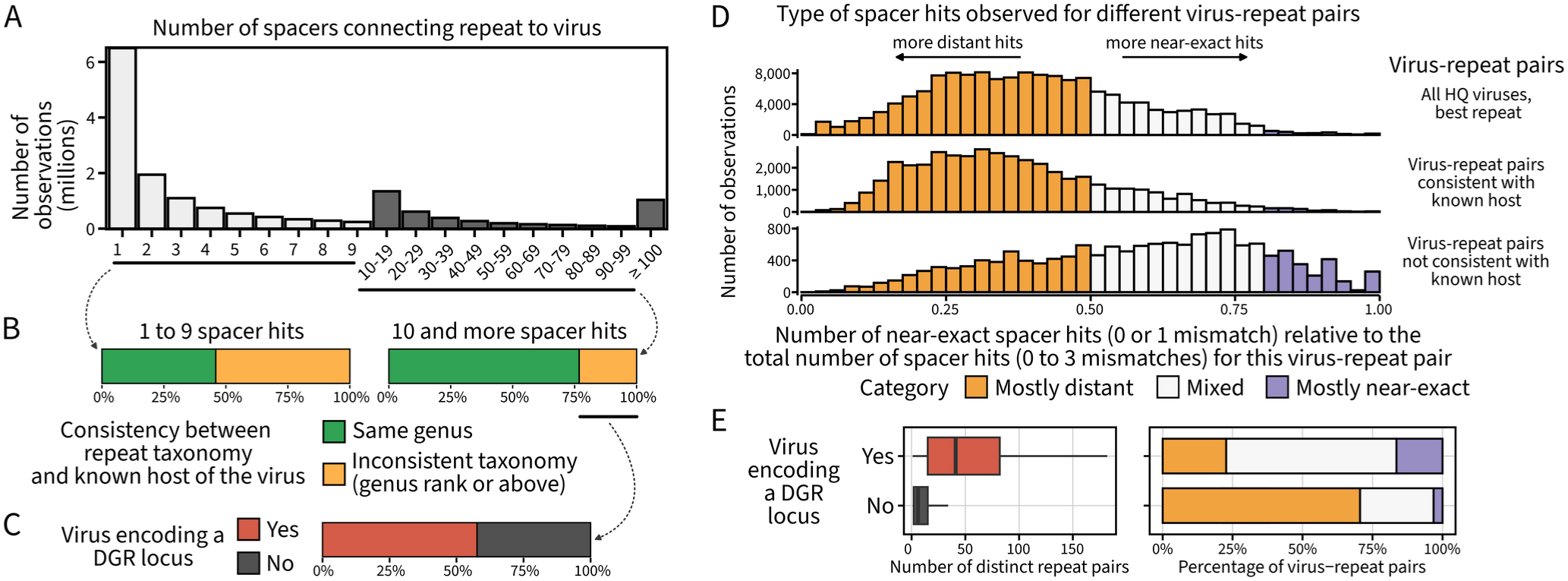
Coverage patterns of potential targets by CRISPR spacers. **A.** Distribution of the number of spacers connecting individual CRISPR repeats to individual IMG/VR virus sequences. Virus-repeat pairs connected by more than 10 spacers are highlighted in black. **B.** Consistency between the taxonomic assignment of the repeat and the known host of the virus, inferred from detection as a prophage/provirus or as an isolated virus. The proportion of consistent and inconsistent taxonomies is presented for virus-repeat pairs connected by 1 to 9 spacer hits, and virus-repeat pairs connected by 10 and more spacer hits. **C.** Percentage of high-quality viruses predicted to encode a DGR locus, for viruses involved in a virus-repeat pair with ≥10 spacer hits and inconsistent taxonomy between the repeat and the known virus host. **D.** Distribution of the ratio of near-exact spacer hits (0 or 1 mismatch) over total number of spacer hits (0 to 3 mismatches), for individual virus-repeat pairs. From top to bottom, this distribution is shown for (i) high-quality viruses and the repeat with the highest number of spacer hits for each, (ii) high-quality viruses with known hosts, and all repeats with a taxonomic assignment consistent with this known host, and (iii) high-quality viruses with known hosts, and all repeats with a taxonomic assignment inconsistent with this known host. **E.** Relationship between DGR detection, number of repeats with ≥10 spacer hits, and mismatch profile category based on the ratio of near-exact spacer hits to the total number of spacer hits (see panel D), for high-quality viruses. For the boxplot in the left panel, the box shows the inter-quartile range (IQR), with the median indicated by a vertical bar, and whiskers extend to 1.5*IQR below the first quartile and above the third quartile.

We next explored the 23.2% of virus-repeat pairs that included ≥10 near-exact spacer hits but showed a discrepancy between the targeting taxon and the known host of the virus (Fig. 4B). Strikingly, 57.6% of these cases involved viruses encoding a Diversity-Generating Retroelement (DGR), compared to 16.8% across all pairs. DGRs are targeted mutagenesis mechanisms that, in phages, can modify receptor binding proteins and lead to virion attachment to non-host cells^48^. Previous smaller scale studies already highlighted that DGR-encoding viruses are targeted by a broader diversity of CRISPR systems than non-DGR-encoding viruses^49^, and the large spacer database collected here further suggests that some of these may originate from non-host microbes.

To further understand whether CRISPR targeting may be observed, in some cases, for non-host microbes, we selected all HQ virus-repeat pairs with ≥10 near-exact spacer hits, and tallied the number of more distant spacer hits (2 or 3 mismatches between spacer and virus) for these same pairs. For known phages, expected virus-host pairs were dominated by distant spacer hits (Fig. 4D, Fig. S13). This high number of distant hits for known hosts is consistent with compensatory mutations being selected over time to avoid CRISPR targeting^48–50^. Meanwhile, repeats assigned to a different (non-host) microbial taxon typically showed a majority of near-exact hits (Fig. 4D, Fig. S13). These CRISPR targeting may thus be linked to phage genome entry into a non-host cell, from which no compensatory mutations are expected. When considering the best (i.e. highest number of spacer hits) pair for all HQ viruses, the majority indeed showed a majority of distant hits, and would be consistent with virus-host interactions (Fig. 4D). However, DGR-encoding viruses seemed to be frequently associated with a larger number of repeats, including many for which the spacer hits were mostly near-identical (Fig. 4E).

Taken together, these observations are consistent with DGRs enabling attachment of virions to non-host cells, after which the CRISPR system(s) of this non-host cell may acquire spacers following virus genome entry. Contrary to CRISPR targeting by a host cell, no selection for escape mutants would be expected as other defense systems and/or a lack of adaptation to the cellular machinery would likely prevent successful infection regardless. This scenario would explain the different ratio of near-exact spacer hits observed for DGR-encoding viruses, and would also apply to other cases in which a virus genome would repeatedly enter a non-host cell. For instance, cell-to-cell transfer (e.g. conjugation or inter-species mating^53^), or natural transformation following lysis of a neighboring cell, could lead to recurring interactions between viruses and specific non-host microbes, especially in structured microbiomes with limited local diversity such as biofilms, micro-aggregates, or between syntrophic partners^38^. Ultimately, these data suggest that the number of unique spacer hits and the number of mismatches between spacer and virus are relevant metrics when trying to infer virus-host interactions from CRISPR targeting.

### Broad CRISPR targeting is relatively rare and does not necessarily reflect broad host range

Finally, in light of this expanded CRISPR spacer database and different types of virus-microbe interactions identified, we sought to analyze in more detail viruses seemingly targeted by multiple unrelated taxa. Several such cases were recently reported and may suggest the existence of viruses with a much broader host range than any experimentally characterized isolate, possibly spanning across multiple classes, phyla, or even domains^37–39,50^. However, most of these observations were based on a limited number of spacer matches and could also derive from sequences shared across unrelated viruses or from the presence of a viral genome in a non-host cell^38^.

First, several viruses from human and landfill samples previously described as targeted by distinct phyla^37,39^ also appeared as broadly targeted in our analysis (Fig. 5A). Most of these viruses showed a “dominant” targeting pattern, i.e. one taxon encoding spacers covering a large fraction of the virus genome (≥25% and ≥10kb), while the targeting by other taxa was limited to <300bp (Fig. S15). One exception was the landfill virus I49 which showed relatively large targeting (1kb and 900bp) by two distinct phyla (*Muirbacteria* & *Patescibacteria*), with spacer hits evenly distributed across the ∼20kb of the virus sequence, but the corresponding mismatch profiles did not clearly reflect a likely virus-host interaction (Fig. 5A, Fig. S15, Table S3, Supplementary Text 5). In addition to previously reported sequences, several additional viruses could be identified in the same landfill datasets, including two (I15 and I10) showing broad spacer coverage (≥1kb) by distantly related taxa ( *Bacilli* & *Negativicutes*, and *Halobacteriota* & *Clostridia*) with a mismatch profile consistent with virus-host interactions, including one (I15) encoding a DGR (Fig. 5B, Fig. S16, Table S3, Supplementary Text 5).

**Fig. 5.**
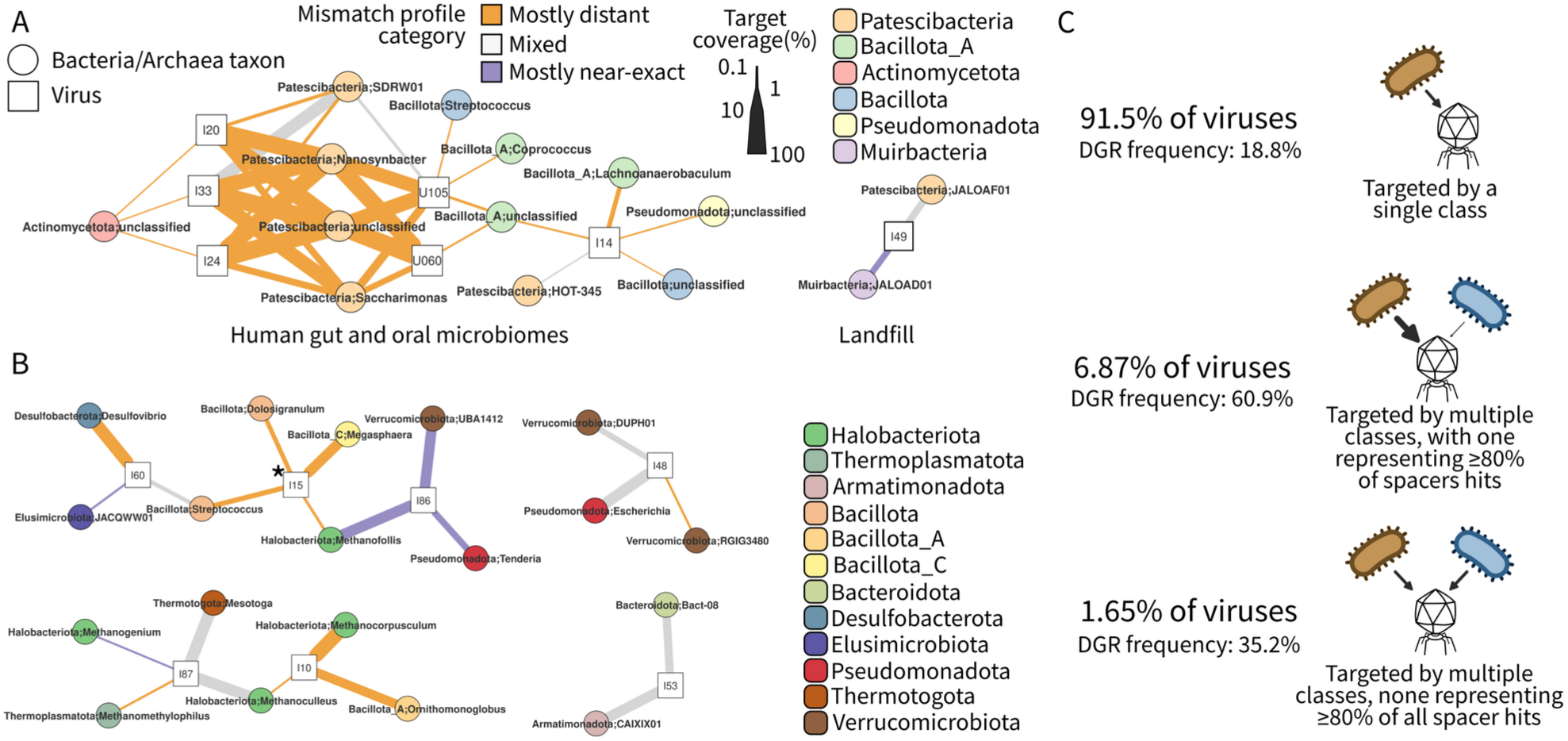
CRISPR targeting of viruses across microbial taxa. **A.** Virus-taxon network for viruses identified as targeted by multiple phyla in previous studies^37,39^ and in this analysis. Microbial taxa are grouped at the class ranks and represented by circle nodes colored by phylum and labeled with the corresponding microbial class. Viral contigs are represented by square nodes, and labeled with a simplified IMG/VR or UGV id (I20: IMGVR_UViG_3300008436_000020, I33: IMGVR_UViG_3300008506_000033, I24: IMGVR_UViG_3300008155_000024, U105: UGV-GENOME-0183105, U060: UGV-GENOME-0152060, I14: IMGVR_UViG_7000000432_000014, I49: IMGVR_UViG_3300028603_000349). Connections between viruses and taxa are based on all spacers from this taxon matching this virus, with the edge width scaled by the target coverage, and the edge colored based on the mismatch profile (see Fig. 4D). **B.** Virus-taxon network for virus sequences assembled from landfill metagenomes ^37^ and connected to multiple phyla. The network displays microbial taxa (circle nodes), virus sequences (square nodes), and connections based on spacer hits (edges) using the same format as panel C. Virus sequence identifiers are simplified IMG/VR ids (I60: IMGVR_UViG_3300028601_000060, I15: IMGVR_UViG_3300014203_000115, I86: IMGVR_UViG_3300028603_000086, I48: IMGVR_UViG_3300014206_000048, I87: IMGVR_UViG_3300028602_002887, I10: IMGVR_UViG_3300028601_000010, I53: IMGVR_UViG_3300029288_007053). The asterisk symbol for virus I15 indicates that this virus genome encodes a DGR. **C.** Overview of the diversity of repeats targeting individual IMG/VR viruses. Only repeats with a class-level taxonomic assignment or better were included, and only cases in which viruses were targeted by ≥10 spacer hits for a given taxon were considered (n=563,062 viruses). Viruses were divided into 3 groups, from top to bottom: viruses targeted by a single class, viruses targeted by multiple classes with one accounting for ≥80% of the spacer hits, and viruses targeted by multiple classes with none accounting for ≥80% of the spacer hits. The corresponding percentage of viruses in the dataset is indicated next to each category, as well as the percentage of high-quality viruses within this group that encode a DGR.

Expanding this analysis to the entire dataset, we observed that a vast majority (91.5%) of viruses were connected to a single bacterial or archaeal class, when considering virus-taxon pairs with ≥10 spacer hits (Fig. 5C). Pairs of repeats assigned to distinct classes but targeting the same virus were typically co-detected in the same samples, suggesting the microbes encoding these repeats may indeed co-exist in the same environment and be exposed to similar viruses (Fig. S17). For viruses targeted by multiple classes, a majority (80.7%) showed one class representing most of the targeting (≥80% of the spacer hits). This group was especially enriched in DGR-encoding viruses (60.9% of high-quality viruses), consistent with the hypothesis that DGR could enable frequent entry into non-host cells (Fig. 5C). For the remaining 9,277 vOTU, some of the most common targeting taxa included several known to engage in microbe-microbe interactions and cross-feeding such as *Phascolarctobacterium*^51^, and several known to aggregate in multispecies biofilms such as *Megasphaera*^52^, both processes potentially leading to frequent contacts between individual viruses and diverse microbes. While errors in the taxonomic assignment of the repeat and horizontal transfer of arrays across unrelated microbes could lead to similar observations, these cases represent the most likely candidates for viruses with unusually broad host range in our dataset, if such viruses exist.

## Discussion

CRISPR-Cas systems are critical molecular mechanisms for microbes to identify and degrade allochthonous DNA and RNA elements. They are especially important as a defense mechanism against viruses, although counter-defenses including anti-CRISPR proteins and escape mutations can limit their efficiency and lead to arm races dynamics^16,19^. The global-scale database of CRISPR spacers built here from public metagenomes reveals that most CRISPR arrays appear to be active and diverse, especially in hot springs and bioreactor ecosystems, possibly due to highly structured micro-habitats such as heterogeneous biofilms and aggregates existing in these environments. This is in contrast with a recent study suggesting a relatively low rate of new spacer acquisition in some human gut microbes^21^. Differences in methodologies may explain these seemingly contradictory results: given the high diversity of spacers observed here and via deep PCR sequencing^36^, it is likely that CRISPR arrays are indeed highly diverse in natural populations, and CRISPR spacers frequently acquired by individual microbes. However, most of these spacers may remain confined to only a few individuals, and only a small minority may propagate through the host population to become prevalent. Conservative analysis frameworks focusing on these prevalent spacers may thus report a low rate of new spacer acquisition^21^. Similarly, previous reports of viruses with potentially broad host ranges based on CRISPR targeting by distantly related microbes^37,39^ could be further refined here and, while most seem to be associated with a primary and likely single host taxon, some may represent candidates for broad host range viruses to be further characterized. Overall, the incredible diversity of CRISPR spacers collected from metagenomes, along with the multiple patterns of targeting revealed in this analysis, reflect the extraordinary complexity of microbe-MGE interactions. These findings open new avenues for research into the ecological and evolutionary constraints on and consequences of these interactions across microbiomes.

## Resource availability

Raw data are available at https://portal.nersc.gov/dna/microbial/prokpubs/spacer_database_resources/, and from NCBI SRA using accession numbers listed in the file “Supplementary_Data_File_2_samples.tsv”. Access to the spacer database including spacer information and spacer hits against IMG/VR and IMG/PR is documented at https://spacers.jgi.doe.gov. This companion website also includes notebooks and examples illustrating how to explore and interact with the spacer database, as well as direct links to download the entire database. The main tool used to identify CRISPR spacers in metagenomes, SpacerExtractor, is available at https://code.jgi.doe.gov/SRoux/spacerextractor and as a package on bioconda (https://bioconda.github.io/recipes/spacerextractor/README.html). Additional scripts used to conduct the analysis and prepare figures for this study are available at https://github.com/simroux/globalspacers_scripts.

## Acknowledgments

The work conducted by the U.S. Department of Energy Joint Genome Institute (https://ror.org/04xm1d337), a DOE Office of Science User Facility, is supported by the Office of Science of the U.S. Department of Energy operated under Contract No. DE-AC02-05CH11231. Some of this work was supported by the U.S. Department of Energy, Office of Science, Biological and Environmental Research, Early Career Research Program (SR) awarded under UC-DOE Prime Contract DE-AC02-05CH11231. Some of this work was supported by the Canadian National Science and Engineering Research Council Discovery Grant 2016-03686. LAH was supported by a Canada Research Chair. UG was funded by the European Research Council (grant ERC-AdG 787514).

## Methods

### CRISPR repeat database: Source data, CRISPR repeat identification, and dereplication

To build a global database of CRISPR repeat sequences, CRISPR repeats were searched across 383,318 bacterial and archaeal genomes from NCBI GenBank^53^ and in contigs from 33,727 publicly available and unrestricted assembled metagenomes and metatranscriptomes publicly available in the IMG database^54^ (accessed February 2023).

For genomes and metagenomes, Minced version 0.4.2 (https://github.com/ctSkennerton/minced), derived from CRT^55^, was first run with default parameters to identify potential arrays. For each contig with at least one predicted array, CRISPRCasTyper^40^ v1.8.0 was run (option “--prodigal meta”) to identify which of these repeat regions may represent CRISPR-Cas systems. From the CRISPRCasTyper prediction, a custom python script (now available as part of “SE_run_cctyper.py” in SpacerExtractor) was applied to refine the predicted repeat sequence from CRISPRCasTyper, specifically looking for cases where the spacers are near-identical at their 5’ and/or 3’ end(s), and extending the repeat as needed. Finally, arrays with the following characteristics were retained: Trusted “True” by CRISPRCasTyper, repeat_identity ≥80, spacer_identity ≤60, spacer_len_sem ≤2, repeat_len ≥2, and repeat_len ≤55. When the repeats were refined by the custom python scripts, these array features were recalculated except for the “Trusted” value from CRISPRCasTyper. This led to a total of 3,168,155 arrays predicted across genomes and metagenomes.

Repeats were first deduplicated, i.e. clustered at 100% nucleotide identity over 100% of both repeats’ lengths (i.e. identical length), via cd-hit-est^56^ v4.8.1 (“cd-hit-est -d 0 -c 1 -g 0 -aL 1 -aS 1”), which yielded 870,235 unique repeats (from 5,808,315 total). Next, these unique repeats were clustered at 100% identity but without constraint on repeat length (“cd-hit-est -d 0 -c 1”). In this second round of clustering, for each genus, only one representative sequence per repeat cluster was kept if this representative sequence represented more than 75% of the repeats in this cluster for this genus. This was done in order to identify and remove potentially mispredicted repeats, i.e. repeats identical to other repeats predicted in the same genus but lacking (or having additional) bases on the repeat’s edge. The cutoff of at least 75% of the repeat sequences being strictly identical was used to ensure that a clear consensus sequence was available to be used as representative. After this second round of clustering, a total of 689,554 unique repeats were obtained.

To further avoid including repeats (and later spacers) unlikely to be linked to a CRISPR array, the 689,554 unique repeat sequences were further screened for low complexity based on Local Composition Complexity values and kmer frequency profiles. Specifically, we used the lcc_simp function from the Bio.SeqUtils.lcc python package and flagged all repeats with lcc value <0.8 as “low complexity”. In addition, we used the kcounter package to get counts of kmers from k=2 to k=8, and flagged repeats based on the frequency of the majority kmer (≥30% at k=2, ≥20% at k=3, ≥5% at k=4, ≥3% at k=5, and ≥2% at k=6, 7, and 8). This procedure identified 107,628 “low-complexity” repeats that were filtered out. Finally, all metagenome-derived repeats with a taxonomic assignment to *Eukaryota* or to viruses outside of the *Caudoviricetes* order (see below) were considered as more likely to originate from a repeat region not associated with a CRISPR array, and excluded (n= 315). The final curated database of repeats was composed of 581,889 unique repeats (Supplementary Data File 1). These repeat sequences were further compared to level 4 CRISPR preditions from CRISPRCasDb^57^ (April 14, 2022 version), and the corresponding CRISPRCasDb identifiers were included in Supplementary Data File 1 when identified.

### CRISPR repeat database: Taxonomic and CRISPR type assignment process

Taxonomic annotation of the repeats (GTDB taxonomy version r214) was based on genome taxonomy for detections made in a genome or MAG, or on contig taxonomy otherwise. For each unique repeat, the taxonomic assignment was based on the lowest common ancestor (LCA) of all identical repeats detected in isolate genomes if any, or the LCA of MAG contigs larger than 500kb if any, or the LCA of all MAG contigs. Because of challenges with binning contigs encoding repeated structures such as CRISPR loci, we divided genome-based taxonomy into three different confidence tiers based on the number and type of detections as follows: (i) the taxonomic assignment of repeats found in 4 or more genomes of isolates was considered as “high-confidence” (n=3,073), (ii) the taxonomic assignment of repeats found in 4 or more MAGs or 1 or more isolate was considered as “medium-confidence” (n=18,572), and (iii) the taxonomic assignment of repeats found exclusively in MAGs and in less than 4 genomes was considered as “low-confidence” (n=21,838). Repeats exclusively detected in metagenomes (n=538,406) were classified based on the taxonomic assignment of the corresponding contig in IMG (April 2024), or IMG/VR for contigs predicted to be of viral origin, considering only cases in which this contig classification was based on at least 2 assigned genes, and taking the LCA of all classified contigs for repeats identified across multiple datasets. This yielded an additional 66,327 taxonomic assignments of individual repeats to bacterial (n=62,076), archaeal (n=2,987) or phage (*Caudoviricetes*, n=1,264) taxa.

In this process, some repeats were assigned to eukaryote (n=11) or other viruses (n=304) and were removed from the final curated database, as mentioned above. The remaining unique repeats (n=472,079) were not assigned to any taxon. The vast majority of these unclassified repeats (94%, n=442,389) were exclusively detected on short metagenomic contigs with 0 or 1 predicted gene, i.e. the contig consisted almost entirely of the putative CRISPR array without any taxonomically-informative region. Finally, each unique repeat was assigned to a predicted CRISPR type using CRISPRCasTyper^40^ v1.8.0 (minimum cutoff of 0.75).

### Selection and metadata collection for public metagenomes

To identify shotgun metagenomics datasets relevant for CRISPR spacer search, the NCBI SRA database was searched in December 2023 with the following criteria: source field entered as “Metagenomic”; the strategy field was not one of “Amplicon”, “ATAC-seq”, “Bisulfite-Seq”, “ChIP-Seq”, “CLONE”, “CLONEEND”, “CTS”, “Dnase-Hypersensitivity”, “EST”, “FAIRE-seq”, “FINISHING”, “FL-cDNA”, “GBS”, “Hi-C”, “miRNA-Seq”, “ncRNA-Seq”, “POOLCLONE”, “RAD-Seq”, “RIP-Seq”, “RNA-Seq”, “Synthetic-Long-Read”, “Targeted-Capture”, “Tethered Chromatin Conformation Capture”, or “Tn-Seq”; the selection field was one of “other”, “PCR”, “RANDOM”, “RANDOM PCR”, “size fractionation”, or unspecified; and the sequencing platform was one of “BGISEQ” or “ILLUMINA”. Datasets indicated as containing less than 1,000,000 spots or with an average read length lower than 90 in the SRA metadata were excluded, as were datasets that likely underwent whole-genome amplification, such as those mentioning single-cell sorting or MDA, because whole-genome amplification often leads to over-inflated predicted spacer diversity, possibly including erroneous spacers due to amplification errors. Read files from all potential datasets to be mined were then downloaded using Kingfisher^58^, and bbcountunique.sh from BBTools 39.03 was used to evaluate the uniqueness of the 25,000 first reads. If the percentage of unique reads in this subset was <20%, the dataset was excluded as either an amplicon dataset or a very low diversity metagenome. The final list of metagenomes used included 467,018 datasets (Supplementary Data File 2).

The NCBI Entrez API was used to identify the BioSample corresponding to each SRA dataset. Each BioSample was then associated with an ecosystem type using a custom script (“biosample_to_ecosystem.pl”) based on the information provided in the fields DATA_env_broad_scale, DATA_env_local_scale, DATA_env_medium, or DATA_Env_Package, the type of package selected (e.g. “DATA_human gut environmental package”), or keywords in the fields DATA_sample comment, DATA_host, DATA_isolation_source, DATA_Organism, DATA_Descr, or Title. When needed and available, information from the Gold^59^ and the MGNify^60^ databases were used to refine the assignment to ecosystem types. This ecosystem information was then aggregated into a simplified classification with 11 ecosystem types: specifically, starting from the detailed classification, ecosystems were ordered based on the number of samples, repeats, and spacers observed, and all ecosystems with <1,500 samples or <10 million spacers were grouped with another ecosystem into a broader category (Fig. S4). Ultimately, 431,265 datasets (92%) could be associated with one of these 11 simplified ecosystem types, with the majority (59%) identified as human-associated.

### Comparison of the number and taxonomy of CRISPR repeats across ecosystems

To compare the number of CRISPR repeats detected across samples and ecosystems while accounting for differences in sequencing depth, we randomly subsampled 1,000 metagenomes from each ecosystem, except for Aquatic;Non-marine_Saline_and_Alkaline and Birds-associated for which 500 metagenomes were subsampled, and Fish-associated and Fungi-associated for which 50 metagenomes were subsampled, because of a lower overall number of metagenomes for these ecosystems. For each metagenome, the number of distinct repeats for which a spacer set was detected (see below) was divided by the total number of bases sequenced in this metagenome. For each ecosystem, the median number of repeats per Gb of metagenome was then tallied. This procedure was repeated 50 times to obtain a distribution of this median number of repeats per Gb across different subsamples for each ecosystem.

The distribution of taxa encoding CRISPR repeats across ecosystems was based on all repeats for which a genus-level taxonomic assignment could be obtained. For each ecosystem, the total number of repeats from each taxon was tallied, and converted into a frequency by dividing by the total number of repeats with a genus-level assignment observed in this ecosystem. A heatmap was then prepared using the ggplot2^61^ v3.5.1 R package and including all genera observed with a frequency ≥2% and/or in the top 3 highest frequencies for at least one ecosystem. A similar approach was used to calculate the frequency of individual CRISPR types detected across ecosystems.

### Initial detection of potential spacers in metagenomic reads

A new approach was designed to identify spacers in metagenomic short-reads data, inspired from CRASS ^35^ and CasCollect^34^. This approach relies on a new component of the BBTools toolkit (“bbcrisprfinder.sh”), available starting in BBTools 39.03 (https://sourceforge.net/projects/bbmap/). The entire pipeline is available in a single package named “SpacerExtractor” (https://code.jgi.doe.gov/SRoux/spacerextractor). The version used in this study corresponds to SpacerExtractor v0.8.

The first step in the pipeline identified and extracted potentially relevant reads based on the 581,889 non-redundant CRISPR repeats previously identified (see above) using bbduk.sh (using a fasta file of these repeats as “ref” argument, minimum kmer length of 23, options kfilter = t, maskmiddle = f). Selected reads were then quality-trimmed using bbduk.sh (options ktrim=r, k=23, mink=11, hdist=1, tbo, tpe, minlen=90, ref=adapters, ftm=5, maxns=0, maq=8) and merged if sequenced as paired-end with bbmerge.sh (options mix, nn=t). Next, bbcrisprfinder.sh was used to identify potential “repeat – spacer – repeat” loci in these selected and trimmed reads in two ways: one using the 581,889 non-redundant CRISPR repeats as guides (options k=13, mm=1, minrepeats=2, rmismatches=1, minspacer=20, maxspacer=55, minrepeat=20, maxrepeat=55, bruteforce=t, mintailrepeat=6, rmmt=0, kref=13, mmref=0, maxrefmm=0), the other leveraging the *de novo* detection of “repeat – spacer – repeat” loci from bbcrisprfinder.sh (options k=13, mm=1, minrepeats=2, rmismatches=1, minspacer=20, maxspacer=55, minrepeat=20, maxrepeat=55, bruteforce=t, mintailrepeat=6, rmmt=0). For the *de novo* detection, two rounds of detection were performed, with the second one using the repeat sequences identified in the first *de novo* detection as guides. The results of these two different spacer detection steps were then merged to identify the most likely repeat for each read, with priority given to exact complete repeat (2 full-length repeats, no mismatch) over exact partial repeat (1 full-length repeat and 1 partial repeat) and over non-exact repeats (some mismatch between the two copies of the repeat). If multiple instances of the same repeat were detected on the same read, the quality of this repeat in this read was taken as the highest quality across all repeats, and the total number of predicted “repeat-spacer-repeat” regions was tallied.

Eventually, each read was assigned to a single repeat based on (i) number of “repeat-spacer-repeat” regions detected (higher is better), (ii) length of the repeat (higher is better), (iii) quality of the best repeat matches (see above), and (iv) global count of the repeat in the metagenome (higher being better). After linking a read to individual repeats, additional spacers were searched for the same repeat on the read based on the detection of partial repeats on the read’s 5’ and 3’ edges that would have been missed in the first round of detection (minimum of 4bp with no mismatch for considering a partial repeat on a read’s 5’ or 3’ edge). For each metagenome library, this process yielded a list of potential repeats and spacers that were then further refined and quality-checked (Fig. S3).

### Identification of potentially spurious CRISPR spacers

To remove as many spurious spacer detections as possible, potential spacers were first dereplicated for each metagenome library at 100% nucleotide identity using cd-hit-est^56^ v4.8.1 (options -c 1, -G 0, -aL 1, -aS 1). Next, because the repeats and spacers were detected from individual reads and not used in a consensus assembly, a denoising step similar to the one used for 16S rRNA amplicons in DADA2^62^ was performed (Fig. S3). Specifically, the repeats detected in a given library were first denoised to identify cases in which a repeat detection likely derived from a sequencing error based on the presence of a near-identical repeat with much higher coverage in the same library. This was done by calculating an expected number of cases in which a given repeat R1 would be observed instead of another repeat R2 due to sequencing error (assuming an error rate of 0.001), considering only cases in which the overall coverage of R2 is higher than the one of R1. A p-value associated with this expected number of observations under a sequencing error scenario was then calculated based on a Poisson distribution, and a repeat R1 was reclassified as R2 if this p-value was ≥0.05, unless the repeat R1 was a singleton (i.e. observed only in one read) in which case it was reclassified if the number of expected observations was ≥0.05. When reclassifying repeats, all reads on which the original repeat R1 was predicted were then reprocessed to extract new spacer sequences corresponding to R2.

Next, a similar approach was used to denoise the spacer sequences predicted for each repeat. For each predicted repeat R1, spacers were compared to each other and a spacer S1 was reclassified as spacer S2 if it was likely to be derived from S2 given expected sequencing errors (p-value ≥0.05, or expected number of observations ≥0.05 for singleton spacers). Finally, potentially erroneous spacers were flagged after denoising based on the following criteria (Fig. S3): (i) spacer sequences identified as low quality or low complexity, i.e. a Local Composition Complexity <0.8 as computed with the Bio.SeqUtils lcc_simp function, the presence of an ambiguous base (“N”) in the spacer sequence, or high frequency of a given kmer across the spacer sequence suggestive of a repeat region within the spacer; (ii) detection on the edge of a read, i.e. spacers for which one repeat is partial and <5nt; (iii) spacers with an outlier length compared to other spacers associated with the same repeat in this sample, i.e. with a length either shorter than the first quartile minus 1.5*inter-quartile range, or longer than the third quartile plus 1.5*inter-quartile range, based on the length of all spacers for this repeat in this sample; and (iv) spacers with unexpectedly conserved 5’ or 3’ ends. Spacer conservation is based on deduplicated spacers and computed as follows: for terminal 2-, 3-, and 4-mers, spacer ends were considered as unexpectedly conserved if the most frequent k-mer was found in >50% of deduplicated spacers and with a p-value <1E-5 when calculating the probability of observing this number of identical k-mers at random under a binomial distribution given the size of the kmer and the total number of deduplicated kmers observed. If unexpected spacer end conservation was detected, all spacers associated with this repeat in this sample were ignored. For cases in which identical spacers were associated with multiple repeats in the same metagenome library, the spacer was assigned only to the repeat for which the spacer prediction was the highest quality (i.e. no warning flag), or to multiple repeats if the spacer predictions were of identical quality. Finally, spacers flagged as edge, low-complexity or low-quality, or with unexpected length or conservation patterns, and spacers associated with rare repeats, i.e. repeats for which less than 5 spacers in total were detected in the library (ignoring any spacer flagged with one of the previous warnings), were removed from the final set of spacers predicted for each metagenome.

### Evaluation of spacer recovery from simulated metagenomes

To evaluate the efficiency and limits of our spacer extraction pipeline, we used a simulation framework similar to Zhang et al. ^21^. First, 200 high-quality genomes (<20 scaffolds) with at least one predicted CRISPR array were randomly selected from the IMG database^54^. CRISPRCas typer^40^ v1.8.0, default parameters, was used to establish a ground truth of repeats and spacers for these genomes. Next, art_illumina^63^ v2.5.8 was used to simulate reads from these genomes at 3 different coverage levels (-f parameters set at 0.8, 10, and 100x) and 2 different error profiles (“-ss HS25” and “-ss HS25 -qs 3 -qs2 3”). Other parameters were kept constant (“-p -l 150 -m 200 -s 10”). Three different tools were then used on the simulated reads to predict CRISPR spacers: SpacerExtractor v0.8 (options “-m 0” in “filter_hq_spacers”, high-quality spacers only, default parameters otherwise), Crass^35^ v1.0.1 (default parameters) and MetaCRAST^33^ (May 2023 version). For SpacerExtractor, the default repeat database “SE_Db_r214GBIMG_0.9” was used, for MetaCRAST the sequence of repeats predicted by CRISPRCasTyper was provided for each genome, while Crass predicts both repeats and spacers *de novo*. Only repeats present in the SpacerExtractor database were considered in the results, spacers identical (after reverse-complementing if needed) between CRISPRCasTyper and the *de novo* prediction were considered as true positive (TP), spacers predicted for a known repeat but not listed by CRISPRCasTyper were considered as false positive (FP), and spacers listed by CRISPRCasTyper but not predicted were considered as false negative (FN). Recall was calculated as TP divided by TP+FN, false-discovery rate (FDR) as FP divided by FP+TP, and F1 score as 2*TP divided by 2*TP+FP+FN. In addition, another set of metrics was calculated from the *de novo* spacers predicted by SpacerExtractor after ignoring all singleton spacers (coverage=1).

### Clustering of spacers and spacer set diversity

In order to build a non-redundant database of spacers, the 794,125,610 curated spacers were clustered at 100% identity using a custom python script looking for exact string matches (over the entire length) between two spacers or between a spacer and the reverse-complement of another spacer. To evaluate the sequence similarity of singleton spacers to other spacers in the same sample, cd-hit-est v4.8.1 was used to cluster spacers extracted from each individual sample and each repeat at 95% identity (-c 0.95) and 80% identity (-c 0.80).

In order to define spacer sets, a list of all distinct spacers associated with a given repeat was obtained for each sample, and considered as the “spacer set” for this repeat in this sample. Each spacer was associated with an estimated depth of coverage based on the number of reads in which this spacer was identified in the corresponding sample. Based on the observed distribution of spacer relative coverage within sets, two categories of spacers were empirically defined: “common” spacers as spacers with a number of reads ≥50% of the number of reads of the highest coverage spacer (i.e. maximum number of reads) in the set, and “rare” spacers as spacers with a number of reads <5% of the number of reads of the most covered spacer (Fig. S6). At the spacer set level, we categorized spacer sets as “low” diversity if ≥50% of the spacers in the set were common. Spacer turnover between samples was evaluated by identifying and tallying the number of identical spacers between spacer sets connected to the same repeat. This was done only for repeats associated with ≥10 sets of ≥10 spacers or more and at least one spacer with ≥20x coverage (Fig. S6). To account for differences in the number of samples in which each repeat is found, the distribution of individual spacers was estimated for a random subset of 10 samples for each set, and the number of samples in which each spacer was observed was tallied. Spacers were considered as “common” if they were considered as common (coverage ≥50% of the maximum spacer coverage in the corresponding spacer set) in at least one sample.

Samples originating from individual subjects (for host-associated samples) or individual sampling sites (for environmental samples), were identified based on the corresponding BioSample metadata. Specifically, samples with the same value in the “DATA_host_subject_id” field were considered as coming from the same subject, and samples with the same value in the “DATA_lat_lon” field were considered as coming from the same location. Spacer distribution was evaluated using the same approach as previously described: repeats for which spacer sets including ≥10 spacers with at least one spacer with ≥20x coverage were available for ≥10 different sampling dates (to avoid exact replicates) for an individual subject or location were identified. For each of these sets, a random subset of 10 samples from an individual subject of location was selected, and the number of samples in which each spacer was observed was tallied.

### Matching spacers to IMG/VR and IMG/PR sequences

Bowtie1 v1.3.1^64^ was used to identify matches between the 409,584,943 non-redundant spacers and either 5,579,078 IMG/VR v4.1^65^ high-confidence virus genomes or 699,973 IMG/PR v1 plasmids^66^. Specifically, a bowtie1 database was built for IMG/VR v4.1 and IMG/PR v1, and spacers were mapped to each database using Bowtie v1.3.1 with the following options: “ -a -v 3” to report all alignments (“-a”) for a given spacer and report alignments with up to 3 mismatches (“-v 3”) over the whole length of the spacer. The resulting sam file is then parsed to extract the coordinates of each match along with the number of mismatches (from the “NM” tag), the CIGAR string, and the MD string. Individual alignments of spacers to IMG/VR and IMG/PR sequences were further filtered to remove all alignments overlapping with (i) a low complexity region in the target, detected with dustmasker v1.0.0 (package blast 2.5.0)^67^ with default parameters, or (ii) a region predicted as being a CRISPR array, detected with minced v0.4.2 (default parameters; https://github.com/ctSkennerton/minced) and by mapping sequences from the repeat database built in this study using Bowtie v1.3.1.

To estimate the number of hits that would be expected from random sequences of a similar length and nucleotide content, we randomly selected a subset of 1 million spacers with at least one hit in IMG/VR, reversed the sequences, and used the same approach as described above to match to IMG/VR. A total of 309 reversed spacers had a hit to an IMG/VR sequence with 0 or 1 mismatch, suggesting a random hit rate of ∼ 0.03%.

In order to evaluate the connection between CRISPR targeting and virus-host relationships, we identified 233,982 sequences in IMG/VR with “known” hosts, i.e. genomes with a host indicated in the NCBI RefSeq Virus database (n=3,438) or genomes detected as prophages or proviruses in bacterial or archaeal genomes, respectively (n=230,544). For these genomes, we specifically compared the host taxonomy as indicated in NCBI RefSeq Virus or in IMG (for prophages/proviruses) to the taxonomy of the repeat for which spacers targeting this genome were identified, after converting all taxonomies to GTDB r214. The taxonomies were considered as consistent if they were identical from domain to genus rank, and inconsistent if different at any rank, e.g. pointing to a different genus, family, order, etc.

### Prediction of protospacer-adjacent motifs (PAM) based on matches to IMG/VR and IMG/PR

For all spacer hits identified with 0 or 1 mismatch across both IMG/VR and IMG/PR, the 10-bp regions upstream and downstream of the match were extracted. For each repeat cluster, surrounding regions across all spacer hits were deduplicated such that only 1 copy of each region was retained when both upstream and downstream 10-bp sequences were identical. Then, conserved nucleotides observed in ≥75% of the distinct set of regions for a given repeat cluster were identified. These conserved positions were then compared to a set of known protospacer-adjacent motifs (PAMs) previously described in the literature^45,46^ (Fig. S12). Only cases with at least 2 conserved positions were considered, except for repeats predicted as type II for which 1-position motifs were considered based on known PAMs. Because the orientation of the repeat is ambiguous in our analysis, all PAMs were assumed to be upstream of the spacer, except for type II repeats where PAMs were assumed to be downstream of the spacer. If a conserved region was observed on the opposite strand, e.g. upstream of a type II repeat or downstream of a non-type II repeat, the sequence was reverse-complemented for the purpose of PAM detection. Finally, the known PAMs were completed by motifs observed in ≥5% of neighborhoods, which were typically minor variants of known PAMs motifs (Fig. S12).

### Additional annotation, feature prediction, and comparison of IMG/VR genomes

Information on virus genomes including original ecosystem and taxonomic assignment were obtained from the IMG/VR v4 database^65^. In addition, a lifestyle prediction was performed, as well as identification of putative Diversity-Generating Retroelements (DGRs) and anti-crispr genes (Acr). For lifestyle prediction, we identified all sequences for which the IMG annotation included a gene with a product mentioning “integrase” or “recombinase”, and conducted additional searches against a published database of integrase HMM profiles^68^ as well as all HMM profiles annotated as integrase in the PHROG^69^ v4 database (hmmsearch^70^ v3.3.2, score ≥50 and e-value ≤1.0e-05, default parameters otherwise). All sequences predicted to encode an integrase/recombinase based on one or more of these annotations was predicted as having a “temperate” lifestyle. For DGRs, the approach used in https://bitbucket.org/srouxjgi/dgr_scripts/src/master/DGR_identification/^71^ was used, with a detection of putative DGR reverse-transcriptase based on a collection of HMM profiles (hmmsearch^70^ v3.3.2, score ≥50 and e-value ≤1.0e-05, default parameters otherwise) followed by a custom script to detect likely DGRs based on the presence of a nearby TR-VR structure. Finally, anti-CRISPR genes were searched based on the APIS database^72^ (June 2024) using diamond^73^ v2.1.8 (“--max-target-seqs 999999999”, score ≥50, e-value≤1e-05, minimum coverage of 80% for query and target, default parameters otherwise) and hmmsearch^70^ v3.3.2 (minimum score of 50 and maximum e-value 1.0e-05, default parameters otherwise).

To compare the functional annotation of targeted and untargeted regions in viruses, subsamples of 1,000,000 spacers matching a single vOTU and 1,000,000 spacers matching multiple vOTUs were selected. For the corresponding targets, the predicted cds overlapping the spacer hits were extracted from the IMG annotation, and annotated with the PHROG^69^ v4 database using MMseqs2^74^ version 14.7e284 (easy-search function, score ≥50, e-value ≤1e-05, default parameters otherwise). Genes were considered as functionally annotated if they showed one or more hits against the PHROG database.

To evaluate the similarity between sequences targeted by the same spacer, a set of 2,000,000 spacers matching multiple high-quality virus genomes was randomly selected, and genomes targeted by each spacer were compared all-vs-all using lz-ani^75^ v1.0.1 (all2all function, default parameters). For each spacer, the largest distance (i.e. 1 -gani) between a pair of high-quality virus genomes was taken as the maximum distance between targets for this spacer.

### Definition of targeting patterns and identification of associated viral features

For IMG/VR matches, patterns of spacer targeting (i.e. number of spacers matching a target, percentage of target covered, number of spacer hits for different number of mismatches between spacer and target, and targeting by prokaryotic host taxon) were calculated for prokaryotic viruses only. Specifically, only genomes assigned to *Tokiviricetes, Caudoviricetes, Faserviricetes, Malgrandaviricetes, Huolimaviricetes, Leviviricetes, Tectiliviricetes,* or *Laserviricetes* classes were considered for these analyses, except sequences assigned to the *Rowavirales* order (i.e. adenoviruses) that were specifically excluded as this taxon contains eukaryotic viruses within the *Tectiliviricetes* class of bacteriophages and archaeoviruses.

To evaluate the level of targeting of individual viruses by individual CRISPR repeats, we analyzed the collection of spacer hits based on all spacers collected across all samples for a given repeat to a given virus genome. The total virus genome coverage by spacers was calculated as the number of positions to which at least 1 spacer from this repeat had a hit with 0 or 1 mismatch, divided by the total length of the virus genome.

For virus-repeat pairs including ≥10 spacer hits with 0 or 1 mismatch covering ≥200bp and for which the virus genome is of “high-quality” (i.e. ≥90% completeness), the spacer hits with 2 or 3 mismatches were extracted from the same Bowtie1 mapping (see above) and tallied to be compared with the number of spacers matching the same virus genome with 0 or 1 mismatch. To summarize these mismatch profiles we computed the ratio between the number of spacer hits with 0 or 1 mismatch divided by the total number of spacer hits (0, 1, 2, or 3 mismatches). This ratio was then used to empirically define 3 categories of virus-repeat pairs: “mostly distant” if the ratio was <0.5, i.e. a majority of spacer hits are found with 2 or 3 mismatches, “mostly near-exact” if the ratio was ≥0.8, i.e. more than 80% of the spacer hits show 0 or 1 mismatch, or “mixed” for ratios between 0.5 and 0.8 (Fig. 4D).

### Analysis of targeting patterns for viruses with potentially broad host range

When exploring the taxonomic range of CRISPR repeats targeting individual viruses, only repeats with a taxonomic assignment of medium-confidence or high-confidence were used (see “CRISPR repeat database: Taxonomic and CRISPR type assignment process” above). Next, virus sequences from two previous studies^37,39^ that highlighted potential broad range targeting were re-analyzed using this set of taxonomically assigned spacers. The sequences of 547 and 13,404 viruses, respectively, were collected and spacer mapping using the same bowtie1-based approach was performed for the same non-redundant set of spacers (see above). Spacer hits were collated using the same approach as previously described to calculate overall coverage and mismatch profile for each virus-taxon pair.

For virus-taxon network representation, two selection criteria were applied. First, the specific virus sequences that had been previously highlighted as targeted by multiple phyla^37,39^ were selected and included in the virus-taxon network if also connected to multiple phyla based on hits from the global spacer database (panel C of Fig. 5). Next, virus sequences from the George et al. dataset^37^ not already included in panel A but connected to repeats from different phyla with 10 or more hits based on the new global spacer database were included in a separate network (panel B of Fig. 5). In both cases, Cytoscape^76^ 3.10.1 was used to display the networks.

### Visualization and statistical tests

All visualization and statistical tests were performed using the R software, version 4.3.3 (2024-02-29), except for the networks which were displayed with Cytoscape version 3.10.1.

## References

1. Falkowski, P.G., Fenchel, T., and Delong, E.F. (2008). The Microbial Engines That Drive Earth’s Biogeochemical Cycles. Science 320, 1034–1039. 10.1126/science.1153213.

2. Gilbert, J.A., Blaser, M.J., Caporaso, J.G., Jansson, J.K., Lynch, S.V., and Knight, R. (2018). Current understanding of the human microbiome. Nat Med 24, 392–400. 10.1038/nm.4517.

3. Macalady, J.L., Hamilton, T.L., Grettenberger, C.L., Jones, D.S., Tsao, L.E., and Burgos, W.D. (2013). Energy, ecology and the distribution of microbial life. Phil. Trans. R. Soc. B. 10.1098/rstb.2012.0383.

4. West, S.A., Diggle, S.P., Buckling, A., Gardner, A., and Griffin, A.S. (2007). The Social Lives of Microbes. Annu. Rev. Ecol. Evol. Syst. 38, 53–77. 10.1146/annurev.ecolsys.38.091206.095740.

5. Bottery, M.J. (2022). Ecological dynamics of plasmid transfer and persistence in microbial communities. Current Opinion in Microbiology 68, 102152. 10.1016/j.mib.2022.102152.

6. Breitbart, M., Bonnain, C., Malki, K., and Sawaya, N.A. (2018). Phage puppet masters of the marine microbial realm. Nat Microbiol 3, 754–766. 10.1038/s41564-018-0166-y.

7. Shkoporov, A.N., and Hill, C. (2019). Bacteriophages of the Human Gut: The “Known Unknown” of the Microbiome. Cell Host & Microbe 25, 195–209. 10.1016/j.chom.2019.01.017.

8. Williamson, K.E., Fuhrmann, J.J., Wommack, K.E., and Radosevich, M. (2017). Viruses in Soil Ecosystems: An Unknown Quantity Within an Unexplored Territory. Annu. Rev. Virol. 4, 201–219. 10.1146/annurev-virology-101416-041639.

9. Makarova, K.S., Wolf, Y.I., Iranzo, J., Shmakov, S.A., Alkhnbashi, O.S., Brouns, S.J.J., Charpentier, E., Cheng, D., Haft, D.H., Horvath, P., et al. (2020). Evolutionary classification of CRISPR–Cas systems: a burst of class 2 and derived variants. Nat Rev Microbiol 18, 67–83. 10.1038/s41579-019-0299-x.

10. Barrangou, R., and Horvath, P. (2017). A decade of discovery: CRISPR functions and applications. Nat Microbiol 2, 1– 9. 10.1038/nmicrobiol.2017.92.

11. Doudna, J.A., and Charpentier, E. (2014). The new frontier of genome engineering with CRISPR-Cas9. Science 346, 1258096. 10.1126/science.1258096.

12. Wang, J.Y., and Doudna, J.A. (2023). CRISPR technology: A decade of genome editing is only the beginning. Science 379, eadd8643. 10.1126/science.add8643.

13. Amitai, G., and Sorek, R. (2016). CRISPR–Cas adaptation: insights into the mechanism of action. Nat Rev Microbiol 14, 67–76. 10.1038/nrmicro.2015.14.

14. Barrangou, R., Fremaux, C., Deveau, H., Richards, M., Boyaval, P., Moineau, S., Romero, D.A., and Horvath, P. (2007). CRISPR Provides Acquired Resistance Against Viruses in Prokaryotes. Science 315, 1709–1712. 10.1126/science.1138140.

15. Deveau, H., Garneau, J.E., and Moineau, S. (2010). CRISPR/Cas System and Its Role in Phage-Bacteria Interactions. Annu. Rev. Microbiol. 64, 475–493. 10.1146/annurev.micro.112408.134123.

16. Westra, E.R., Dowling, A.J., Broniewski, J.M., and Houte, S. van (2016). Evolution and Ecology of CRISPR. Annu. Rev. Ecol. Evol. Syst. 47, 307–331. 10.1146/annurev-ecolsys-121415-032428.

17. Fehrenbach, A., Mitrofanov, A., Alkhnbashi, O.S., Backofen, R., and Baumdicker, F. (2024). SpacerPlacer: ancestral reconstruction of CRISPR arrays reveals the evolutionary dynamics of spacer deletions. Nucleic Acids Research 52, 10862–10878. 10.1093/nar/gkae772.

18. Garrett, S.C. (2021). Pruning and Tending Immune Memories: Spacer Dynamics in the CRISPR Array. Front. Microbiol. 12. 10.3389/fmicb.2021.664299.

19. Pawluk, A., Davidson, A.R., and Maxwell, K.L. (2018). Anti-CRISPR: discovery, mechanism and function. Nat Rev Microbiol 16, 12–17. 10.1038/nrmicro.2017.120.

20. Watson, B.N.J., Steens, J.A., Staals, R.H.J., Westra, E.R., and van Houte, S. (2021). Coevolution between bacterial CRISPR-Cas systems and their bacteriophages. Cell Host & Microbe 29, 715–725. 10.1016/j.chom.2021.03.018.

21. Zhang, A.-N., Gaston, J.M., Cárdenas, P., Zhao, S., Gu, X., and Alm, E.J. (2024). CRISPR-Cas spacer acquisition is a rare event in human gut microbiome. Cell Genomics 0. 10.1016/j.xgen.2024.100725.

22. Andersson, A.F., and Banfield, J.F. (2008). Virus Population Dynamics and Acquired Virus Resistance in Natural Microbial Communities. Science 320, 1047–1050. 10.1126/science.1157358.

23. Burstein, D., Sun, C.L., Brown, C.T., Sharon, I., Anantharaman, K., Probst, A.J., Thomas, B.C., and Banfield, J.F. (2016). Major bacterial lineages are essentially devoid of CRISPR-Cas viral defence systems. Nat Commun 7, 10613. 10.1038/ncomms10613.

24. Emerson, J.B., Andrade, K., Thomas, B.C., Norman, A., Allen, E.E., Heidelberg, K.B., and Banfield, J.F. (2013). Virus-Host and CRISPR Dynamics in Archaea-Dominated Hypersaline Lake Tyrrell, Victoria, Australia. Archaea 2013. 10.1155/2013/370871.

25. England, W.E., Kim, T., and Whitaker, R.J. (2018). Metapopulation Structure of CRISPR-Cas Immunity in Pseudomonas aeruginosa and Its Viruses. mSystems 3, 10.1128/msystems.00075-18. 10.1128/msystems.00075-18.

26. Sun, C.L., Thomas, B.C., Barrangou, R., and Banfield, J.F. (2016). Metagenomic reconstructions of bacterial CRISPR loci constrain population histories. ISME J 10, 858–870. 10.1038/ismej.2015.162.

27. Amundson, K.K., Roux, S., Shelton, J.L., and Wilkins, M.J. (2023). Long-term CRISPR locus dynamics and stable host-virus co-existence in subsurface fractured shales. Current Biology 33, 3125–3135.e4. 10.1016/j.cub.2023.06.033.

28. Emerson, J.B., Roux, S., Brum, J.R., Bolduc, B., Woodcroft, B.J., Jang, H.B., Singleton, C.M., Solden, L.M., Naas, A.E., Boyd, J.A., et al. (2018). Host-linked soil viral ecology along a permafrost thaw gradient. Nat Microbiol 3, 870–880. 10.1038/s41564-018-0190-y.

29. Kosmopoulos, J.C., Campbell, D.E., Whitaker, R.J., and Wilbanks, E.G. (2023). Horizontal Gene Transfer and CRISPR Targeting Drive Phage-Bacterial Host Interactions and Coevolution in “Pink Berry” Marine Microbial Aggregates. Appl Environ Microbiol 89, e00177–23. 10.1128/aem.00177-23.

30. López-Beltrán, A., Botelho, J., and Iranzo, J. (2024). Dynamics of CRISPR-mediated virus–host interactions in the human gut microbiome. ISME J 18, wrae134. 10.1093/ismejo/wrae134.

31. Nelson, A.R., Narrowe, A.B., Rhoades, C.C., Fegel, T.S., Daly, R.A., Roth, H.K., Chu, R.K., Amundson, K.K., Young, R.B., Steindorff, A.S., et al. (2022). Wildfire-dependent changes in soil microbiome diversity and function. Nat Microbiol 7, 1419–1430. 10.1038/s41564-022-01203-y.

32. Lei, J., and Sun, Y. (2016). Assemble CRISPRs from metagenomic sequencing data. Bioinformatics 32, i520–i528. 10.1093/bioinformatics/btw456.

33. Moller, A.G., and Liang, C. (2017). MetaCRAST: reference-guided extraction of CRISPR spacers from unassembled metagenomes. PeerJ 5, e3788. 10.7717/peerj.3788.

34. Podlevsky, J.D., Hudson, C.M., Timlin, J.A., and Williams, K.P. (2020). CasCollect: Targeted assembly of CRISPR-associated operons from high-throughput sequencing data. NAR Genomics and Bioinformatics 2, 1–10. 10.1093/nargab/lqaa063.

35. Skennerton, C.T., Imelfort, M., and Tyson, G.W. (2013). Crass: identification and reconstruction of CRISPR from unassembled metagenomic data. Nucleic acids research 41, e105. 10.1093/nar/gkt183.

36. Silas, S., Makarova, K.S.K.S., Shmakov, S., Páez-Espino, D., Mohr, G., Liu, Y., Davison, M., Roux, S., Krishnamurthy, S.R.S.R., Fu, B.X.H.B.X.H., et al. (2017). On the origin of reverse transcriptase-using CRISPR-Cas systems and their hyperdiverse, enigmatic spacer repertoires. mBio 8. 10.1128/mBio.00897-17.

37. George, N.A., and Hug, L.A. (2023). CRISPR-resolved virus-host interactions in a municipal landfill include non-specific viruses, hyper-targeted viral populations, and interviral conflicts. Sci Rep 13, 5611. 10.1038/s41598-023-32078-6.

38. Hwang, Y., Roux, S., Coclet, C., Krause, S.J.E., and Girguis, P.R. (2023). Viruses interact with hosts that span distantly related microbial domains in dense hydrothermal mats. Nat Microbiol 8, 946–957. 10.1038/s41564-023-01347-5.

39. Liu, J., Jaffe, A.L., Chen, L., Bor, B., and Banfield, J.F. (2023). Host translation machinery is not a barrier to phages that interact with both CPR and non-CPR bacteria. mBio 14, e01766–23. 10.1128/mbio.01766-23.

40. Russel, J., Pinilla-Redondo, R., Mayo-Muñoz, D., Shah, S.A., and Sørensen, S.J. (2020). CRISPRCasTyper: Automated Identification, Annotation, and Classification of CRISPR-Cas Loci. The CRISPR Journal 3, 462–469. 10.1089/crispr.2020.0059.

41. Arita, M., Karsch-Mizrachi, I., Cochrane, G., and on behalf of the International Nucleotide Sequence Database Collaboration (2021). The international nucleotide sequence database collaboration. Nucleic Acids Research 49, D121–D124. 10.1093/nar/gkaa967.

42. Daly, R.A., Roux, S., Borton, M.A., Morgan, D.M., Johnston, M.D., Booker, A.E., Hoyt, D.W., Meulia, T., Wolfe, R.A., Hanson, A.J., et al. (2019). Viruses control dominant bacteria colonizing the terrestrial deep biosphere after hydraulic fracturing. Nat Microbiol 4, 352–361. 10.1038/s41564-018-0312-6.

43. Mehta, R.S., Abu-Ali, G.S., Drew, D.A., Lloyd-Price, J., Subramanian, A., Lochhead, P., Joshi, A.D., Ivey, K.L., Khalili, H., Brown, G.T., et al. (2018). Stability of the human faecal microbiome in a cohort of adult men. Nat Microbiol 3, 347–355. 10.1038/s41564-017-0096-0.

44. Mick, E., Stern, A., and Sorek, R. (2013). Holding a grudge: Persisting anti-phage CRISPR immunity in multiple human gut microbiomes. RNA Biology 10, 900–906. 10.4161/rna.23929.

45. Leenay, R.T., Maksimchuk, K.R., Slotkowski, R.A., Agrawal, R.N., Gomaa, A.A., Briner, A.E., Barrangou, R., and Beisel, C.L. (2016). Identifying and Visualizing Functional PAM Diversity across CRISPR-Cas Systems. Molecular Cell 62, 137–147. 10.1016/j.molcel.2016.02.031.

46. Rybnicky, G.A., Fackler, N.A., Karim, A.S., Köpke, M., and Jewett, M.C. (2022). Spacer2PAM: A computational framework to guide experimental determination of functional CRISPR-Cas system PAM sequences. Nucleic Acids Research 50, 3523–3534. 10.1093/nar/gkac142.

47. Neri, U., Camargo, A.P., Bushnell, B., and Roux, S. (2025). Tool Choice drastically Impacts CRISPR Spacer-Protospacer Detection. Preprint at bioRxiv, 10.1101/2025.05.06.652306 https://doi.org/10.1101/2025.05.06.652306.

48. Macadangdang, B.R., Makanani, S.K., and Miller, J.F. (2022). Accelerated Evolution by Diversity-Generating Retroelements. Annu. Rev. Microbiol. 76, 389–411. 10.1146/annurev-micro-030322-040423.

49. Nayfach, S., Páez-Espino, D., Call, L., Low, S.J., Sberro, H., Ivanova, N.N., Proal, A.D., Fischbach, M.A., Bhatt, A.S., Hugenholtz, P., et al. (2021). Metagenomic compendium of 189,680 DNA viruses from the human gut microbiome. Nat Microbiol 6, 960–970. 10.1038/s41564-021-00928-6.

50. Páez-Espino, D., Eloe-Fadrosh, E.A., Pavlopoulos, G.A., Thomas, A.D., Huntemann, M., Mikhailova, N., Rubin, E., Ivanova, N.N., and Kyrpides, N.C. (2016). Uncovering Earth’s virome. Nature 536, 425–430. 10.1038/nature19094.

51. Ikeyama, N., Murakami, T., Toyoda, A., Mori, H., Iino, T., Ohkuma, M., and Sakamoto, M. (2020). Microbial interaction between the succinate-utilizing bacterium Phascolarctobacterium faecium and the gut commensal Bacteroides thetaiotaomicron. MicrobiologyOpen 9, e1111. 10.1002/mbo3.1111.

52. Nallabelli, N., Patil, P.P., Pal, V.K., Singh, N., Jain, A., Patil, P.B., Grover, V., and Korpole, S. (2016). Biochemical and genome sequence analyses of Megasphaera sp. strain DISK18 from dental plaque of a healthy individual reveals commensal lifestyle. Sci Rep 6, 33665. 10.1038/srep33665.

53. Sayers, E.W., Cavanaugh, M., Clark, K., Pruitt, K.D., Sherry, S.T., Yankie, L., and Karsch-Mizrachi, I. (2024). GenBank 2024 Update. Nucleic Acids Research 52, D134–D137. 10.1093/nar/gkad903.

54. Chen, I.-M.A., Chu, K., Palaniappan, K., Ratner, A., Huang, J., Huntemann, M., Hajek, P., Ritter, S.J., Webb, C., Wu, D., et al. (2023). The IMG/M data management and analysis system v.7: content updates and new features. Nucleic Acids Research 51, D723–D732. 10.1093/nar/gkac976.

55. Bland, C., Ramsey, T.L., Sabree, F., Lowe, M., Brown, K., Kyrpides, N.C., and Hugenholtz, P. (2007). CRISPR Recognition Tool (CRT): a tool for automatic detection of clustered regularly interspaced palindromic repeats. BMC Bioinformatics 8, 209. 10.1186/1471-2105-8-209.

56. Fu, L., Niu, B., Zhu, Z., Wu, S., and Li, W. (2012). CD-HIT: accelerated for clustering the next-generation sequencing data. Bioinformatics 28, 3150–3152. 10.1093/bioinformatics/bts565.

57. Pourcel, C., Touchon, M., Villeriot, N., Vernadet, J.-P., Couvin, D., Toffano-Nioche, C., and Vergnaud, G. (2020). CRISPRCasdb a successor of CRISPRdb containing CRISPR arrays and cas genes from complete genome sequences, and tools to download and query lists of repeats and spacers. Nucleic Acids Res 48, D535–D544. 10.1093/nar/gkz915.

58. Woodcroft, B.J., Cunningham, M., Gans, J.D., Bolduc, B.B., and Hodgkins, S.B. (2024). Kingfisher: A utility for procurement of public sequencing data. Version v0.4.0 (Zenodo). 10.5281/zenodo.10525086 https://doi.org/10.5281/zenodo.10525086.

59. Mukherjee, S., Stamatis, D., Li, C.T., Ovchinnikova, G., Bertsch, J., Sundaramurthi, J.C., Kandimalla, M., Nicolopoulos, P.A., Favognano, A., Chen, I.-M.A., et al. (2023). Twenty-five years of Genomes OnLine Database (GOLD): data updates and new features in v.9. Nucleic Acids Research 51, D957–D963. 10.1093/nar/gkac974.

60. Richardson, L., Allen, B., Baldi, G., Beracochea, M., Bileschi, M.L., Burdett, T., Burgin, J., Caballero-Pérez, J., Cochrane, G., Colwell, L.J., et al. (2022). MGnify: the microbiome sequence data analysis resource in 2023. Nucleic Acids Research 51, D753–D759. 10.1093/nar/gkac1080.

61. Wickham, H. (2016). ggplot2: Elegant Graphics for Data Analysis (Springer-Verlag New York).

62. Callahan, B.J., McMurdie, P.J., Rosen, M.J., Han, A.W., Johnson, A.J.A., and Holmes, S.P. (2016). DADA2: High-resolution sample inference from Illumina amplicon data. Nat Methods 13, 581–583. 10.1038/nmeth.3869.

63. Huang, W., Li, L., Myers, J.R., and Marth, G.T. (2012). ART: a next-generation sequencing read simulator. Bioinformatics 28, 593–594. 10.1093/bioinformatics/btr708.

64. Langmead, B., Trapnell, C., Pop, M., and Salzberg, S.L. (2009). Ultrafast and memory-efficient alignment of short DNA sequences to the human genome. Genome Biol 10, R25. 10.1186/gb-2009-10-3-r25.

65. Camargo, A.P., Nayfach, S., Chen, I.-M.A., Palaniappan, K., Ratner, A., Chu, K., Ritter, S.J., Reddy, T.B.K., Mukherjee, S., Schulz, F., et al. (2023). IMG/VR v4: an expanded database of uncultivated virus genomes within a framework of extensive functional, taxonomic, and ecological metadata. Nucleic Acids Research 51, D733–D743. 10.1093/nar/gkac1037.

66. Camargo, A.P., Call, L., Roux, S., Nayfach, S., Huntemann, M., Palaniappan, K., Ratner, A., Chu, K., Mukherjeep, S., Reddy, T.B.K., et al. (2024). IMG/PR: a database of plasmids from genomes and metagenomes with rich annotations and metadata. Nucleic Acids Research 52, D164–D173. 10.1093/nar/gkad964.

67. Camacho, C., Coulouris, G., Avagyan, V., Ma, N., Papadopoulos, J., Bealer, K., and Madden, T.L. (2009). BLAST+: architecture and applications. BMC Bioinformatics 10, 421. 10.1186/1471-2105-10-421.

68. Khedkar, S., Smyshlyaev, G., Letunic, I., Maistrenko, O.M., Coelho, L.P., Orakov, A., Forslund, S.K., Hildebrand, F., Luetge, M., Schmidt, T.S.B., et al. (2022). Landscape of mobile genetic elements and their antibiotic resistance cargo in prokaryotic genomes. Nucleic Acids Research 50, 3155–3168. 10.1093/nar/gkac163.

69. Terzian, P., Olo Ndela, E., Galiez, C., Lossouarn, J., Pérez Bucio, R.E., Mom, R., Toussaint, A., Petit, M.-A., and Enault, F. (2021). PHROG: families of prokaryotic virus proteins clustered using remote homology. NAR Genomics and Bioinformatics 3, lqab067. 10.1093/nargab/lqab067.

70. Eddy, S.R. (2011). Accelerated Profile HMM Searches. PLoS Comput Biol 7, e1002195. 10.1371/journal.pcbi.1002195.

71. Roux, S., Paul, B.G., Bagby, S.C., Nayfach, S., Allen, M.A., Attwood, G., Cavicchioli, R., Chistoserdova, L., Gruninger, R.J., Hallam, S.J., et al. (2021). Ecology and molecular targets of hypermutation in the global microbiome. Nat Commun 12, 1–12. 10.1038/s41467-021-23402-7.

72. Yan, Y., Zheng, J., Zhang, X., and Yin, Y. (2024). dbAPIS: a database of anti-prokaryotic immune system genes. Nucleic Acids Research 52, D419–D425. 10.1093/nar/gkad932.

73. Buchfink, B., Xie, C., and Huson, D.H. (2015). Fast and sensitive protein alignment using DIAMOND. Nat Methods 12, 59–60. 10.1038/nmeth.3176.

74. Steinegger, M., and Söding, J. (2017). MMseqs2 enables sensitive protein sequence searching for the analysis of massive data sets. Nat Biotechnol 35, 1026–1028. 10.1038/nbt.3988.

75. Zielezinski, A., Gudyś, A., Barylski, J., Siminski, K., Rozwalak, P., Dutilh, B.E., and Deorowicz, S. (2025). Ultrafast and accurate sequence alignment and clustering of viral genomes. Nat Methods, 1–4. 10.1038/s41592-025-02701-7.

76. Demchak, B., Hull, T., Reich, M., Liefeld, T., Smoot, M., Ideker, T., and Mesirov, J.P. (2014). Cytoscape: the network visualization tool for GenomeSpace workflows. F1000Res 3, 151. 10.12688/f1000research.4492.2.

