## Supplementary Material for "Planetary-scale metagenomic search reveals new patterns of CRISPR targeting"

**Supplementary Text**

*1. Comparison of approaches for the identification of CRISPR spacers from metagenomic reads*

Because of the repetitive nature of CRISPR array loci, including the presence of near-identical spacers in the same array and the presence of distinct arrays with near-identical repeats on the same genome or in the same microbiome, robustly identifying the set of spacers associated with a given repeat is challenging. Both assembly-based and read-based methods can typically only recover a portion of all spacers encoded, and read-based methods in particular can yield artefactual spacers due to sequencing errors. To illustrate the efficiency of these methods across different levels of sequencing error and genome coverage, we compared our approach to the state of the art programs in the field, CRASS[1] and MetaCRAST[2], using simulated reads from isolate genomes from which CRISPR arrays were previously predicted using CRISPR-Cas Typer[3], as done in ref. [4].

Overall, this benchmarking recapitulated expected trends and highlighted some limitations of CRISPR spacer recovery from metagenomic reads (Fig. S2). Across all tools, both recall (percentage of expected spacers recovered) and false-discovery rate (FDR, percentage of unexpected spacers recovered among all predicted spacers) increased with coverage, i.e., as more reads are available for a given genome, all tools recover more of the original spacers, along with additional erroneous spacers (Fig. S2). The increase in FDR was more important at 100x compared to 10x, and when simulating reads using higher sequencing error rates (HiSeq 2500, vs ~ half of the error of HiSeq 2500 as expected for more recent Illumina sequencing technologies such as NovaSeq). When considering both recall and FDR (F1-measure), MetaCRAST and SpacerExtractor showed similar performances for both error profiles at 10x and 100x, while SpacerExtractor performed better for low-coverage genomes (0.8x) with both a higher recall and lower FDR (Fig. S2). The FDR remained relatively high for genomes with high coverage (100x) and high sequencing error rate despite the denoising procedure included in SpacerExtractor. Ignoring singleton spacers substantially decreased this FDR at high coverage, but at the same time led to a strong reduction in recall for low-coverage genomes (Fig. S2).

This benchmarking thus suggests that (i) all tools only recover a fraction of the CRISPR spacers predicted from the original genome assembly, including at high coverage, and (ii) in some cases (high genome coverage, high sequencing error rate), a substantial number of erroneous spacers can also be predicted, often as singleton spacers. Given that most genomes will be present at relatively low coverage in a metagenome, we opted in this study for using SpacerExtractor with standard denoising and including singleton spacers, however studies focusing specifically on genomes with deep sequencing may want to use a more aggressive denoising procedure and/or remove singleton spacers from their analysis. Importantly, these benchmarks only focus on a set of known repeats, i.e. repeats previously predicted by CRISPR-Cas Typer, and do not take into account the potential of some of these tools to discover of entirely new CRISPR arrays (i.e. new repeats and spacers).

*2. Overall diversity of spacer sets, i.e. diversity of spacers associated with a given repeat in a given sample*

Most spacers sets, defined here as the non-redundant collection of spacers associated with a given repeat in a given sample, and interpreted to reflect the diversity of spacers encoded by members of a given population in this sample, seemed to be composed of a few high-coverage spacers followed by a long tail of rare spacers (Fig. 2B). This pattern may, however, be influenced by singleton spacers, i.e. spacers detected on a single read. These singleton spacers likely include a combination of spacers only acquired and/or encoded by rare population members, spacers having undergone mutation event(s) after acquisition as part of the host genome replication process, and artefactual spacers deriving from errors introduced during the sequencing process. If most of the singleton spacers derive from sequencing errors, the spacer diversity observed in spacer sets may be artificially inflated.

To evaluate the likelihood of singleton spacers deriving mostly from a methodological artifact, we clustered spacers within each spacer set at 95% and 80% identity. We reasoned that “real” rare spacers deriving from independent spacer acquisition events would most likely not be similar to another spacer in the set. Meanwhile, spacers having undergone post-acquisition mutation or deriving from sequencing errors would likely be similar to at least one other spacer in the set, and thus should not be singletons anymore when clustering at lower identity percentage. Across all spacer sets, when clustering spacers at 80% identity within a set, 37.1% of singletons were clustered with another spacer, and may possibly originate from one of the latter two scenarios (spacer mutation after acquisition, sequencing error). The remaining 62.9% likely represent genuine rare spacers (Fig. 2C). This suggests that, overall, CRISPR spacer arrays are most often highly diverse and variable between members of a given microbial population, and that the observed diversity is not primarily explained by artifactual spacers, which is also consistent with previous PCR-based targeted exploration of CRISPR spacers[5].

We next explored if specific ecosystems and/or taxa were preferentially associated with large and diverse spacer sets. After normalizing for coverage depth variations, unusually large spacer sets were most often observed in aquatic biomes, especially thermal springs, and engineered systems. Unusually large spacer sets were also more frequently observed in uncultivated taxa known or predicted to engage in microbe-microbe interactions such as *Candidatus* Kryptonium, *Candidatus* Thiodubiliella, or *Candidatus* Magnetomorum[6–8](Fig. S8). This suggests that specific taxa-environment combinations may select for large and diverse spacer sets, i.e. some populations may be uniquely able to accommodate, and likely benefit from, expanded CRISPR spacer repertoires.

On the other end, a small minority (4.9%) of sets showed a much lower intra-population variation and were instead composed of ≥50% of “common” spacers (Fig. S9). These sets were primarily observed for repeats of type V CRISPR arrays encoded by *Clostridia*, *Bacteroidia*, and *Spirochaetia*, which were frequently associated with a relatively low spacer diversity overall (Fig. S9). These repeats may thus represent arrays with limited rates of spacer acquisition, and for which most spacers may still be retained as “memory” spacers[9]. Further studies of individual CRISPR arrays will however be required to characterize in more detail the eco-evolutionary drivers of spacer acquisition and retention in natural populations[4,10].

*3. Spacer hits to atypical phages and archaeal viruses*

Caudoviruses (tailed phages) are the most common phages and archaeoviruses in IMG/VR and represent the majority of viruses with spacer hits. Beyond caudoviruses however, several other virus taxa also showed sign of CRISPR spacer targeting (Fig. S11). In particular, spacer hits were observed to other dsDNA and ssDNA phages including microviruses (*Malgrandviricetes*) and inoviruses (*Faserviricetes*); atypical archaeal viruses such as bicaudaviruses, fuselloviruses, and globuloviruses; and RNA phages in the levi, cysto, and picobirnaviruses groups. Across all these taxa, the overall patterns of targeting in terms of spacer taxonomy and ecosystem were consistent with expectations. Specifically, dsDNA and ssDNA phages were targeted by a relatively broad diversity of CRISPR spacers found across all types of ecosystems. Conversely, taxa known to infect extremophile archaea were almost exclusively targeted by spacers from archaea-assigned repeats obtained from extreme aquatic environments (Fig. S11). One possible exception was *Aeropyrum* coil-shaped virus (family *Spiraviridae*, IMGVR_UViG_2974665388_000001) for which spacer hits from “marine” samples were obtained (BioSample SAMN06761480), however further examination of the metadata revealed these samples were taken near Vulcano islands, known to harbor extreme archaea including e.g. *Pyrococcus*. Taken together, these results confirm that, overall, spacer hits detected through our global database matching approach seem to represent genuine CRISPR spacer targeting of atypical viruses.

Beyond expected taxa and ecosystems, these spacer hits also revealed potential new hosts for some of these atypical phage and archaeovirus taxa, for which relatively few isolates are available. First, these data confirmed that both microviruses and inoviruses seem to interact with a broad range of microbial taxa, with 193 and 197 distinct families with ≥2 spacer targeting at least one of these ssDNA phage sequences (0 or 1 mismatch), respectively[11,12]. Meanwhile, relatively few spacer hits were identified for known and/or predicted clades of RNA phages, namely leviviruses, cystoviruses, and picobirnaviruses. Specifically, even with the global spacer database, no cystovirus or picobirnavirus could be connected to a CRISPR repeat by more than 1 spacer, and only 48 leviviruses could, out of 82,176 total levivirus sequences in IMG/VR. This low level of CRISPR targeting contrasts with a recent description of a putative new clade of RNA phages, related to partitiviruses, that are highly targeted by a *Roseiflexus* CRISPR array[13]. This may be due to a combination of (i) CRISPR-Cas systems only rarely being used to defend against RNA phages, and/or (ii) the fact that the partiti-like virus targeting was detected based on paired metagenomes and metatranscriptomes sampled from the same site, indicating that RNA phages and CRISPR spacers may diverge too quickly to lead to numerous near-exact matches in a global database search, and targeted sampling may be required to identify these interactions.

*4. Spacer hits to phages and archaeal viruses with known hosts*

To explore the relationship between CRISPR targeting and virus-host interaction, we focused on bacteriophages and archaeal viruses in IMG/VR for which a "known" host is available, either because the virus was isolated and grown on a host culture ("isolate", n=3,438), or because it was detected as part of the whole genome shotgun sequencing of a single bacteria or archaea (summarized as "prophage", n=230,544). The taxonomy (GTDB r214) of these "known hosts" was then compared to the taxonomy assigned to CRISPR repeats (using the same GTDB r214) for which at least one spacer hit was identified to this virus, and 3 cases were identified: (i) both taxonomies were identical from domain to genus rank (ii) taxonomies were inconsistent, e.g. pointing to a different genus, family, or order, and (iii) taxonomies were consistent but one of the two assignments (the "known host" or the CRISPR repeat) was unclassified at the genus rank (and sometimes at higher ranks as well). Cases in this latter category (one of the two taxonomies unclassified at genus rank or higher) were ignored as taxonomy consistency could not be validated in these cases.

For virus-taxon connected by less than 10 spacer hits, 45.9% of cases corresponded to consistent taxonomies. Meanwhile, for virus-taxon connected by 10 spacer hits or more, 76.8% corresponded to consistent taxonomies. This percentage was consistent between "isolate" references (22.5% consistency for <10 spacer hits, and 77.5% consistency for ≥10 spacer hits) and "prophage" references (46.2% consistency for <10 spacer hits, and 76.8% consistency for ≥10 spacer hits).

*5. Targeting patterns for UViGs connected to repeats assigned to multiple phyla*

In previous analyses, several UViGs were highlighted as potentially targeted by repeats assigned to different phyla. Some of these UViGs were also targeted by multiple repeats assigned to different phyla in our analysis, and are displayed in panels C and D of Figure 5. Below is a more detailed description of the targeting patterns observed for these two figure panels.

<u>I24, I20</u>: Spacer hits were detected for multiple repeats assigned to Patescibacteria and *Actinomycetota*, including repeats with high confidence (see Methods) for each phylum. In both cases, the number of spacer hits was low for *Actinomycetota* (≤5), and all hits displayed 2 or 3 mismatches between spacer and (predicted) protospacer.

<u>I33</u>: Spacer hits were detected for multiple repeats assigned to Patescibacteria with high confidence, but only one repeat assigned to *Actinomycetota* with high-confidence. No spacer hits were detected from other repeats assigned to *Actinomycetota* with medium or low confidence. A total of 17 hits were detected between the *Actinomycetota* spacers and this UViG, but only a single hit was detected with 1 mismatch or less.

<u>U105</u>: Spacer hits were detected for multiple repeats assigned to Patescibateria and Bacillota_A, including multiple repeats with high confidence for each phylum. The total number of spacer hits was low for Bacillota_A (≤2 per taxon), and higher for Patescibacteria (up to ~ 3,000 for a single repeat).

<u>U060</u>: Spacer hits were detected for multiple repeats assigned to Patescibateria, Bacillota_A, and *Bacillota*, including multiple repeats with high confidence for each phylum. The total number of spacer hits with 0 or 1

mismatch was low for *Bacillota* and Bacillota_A (≤2 per taxon), and higher for Patescibacteria (up to 363 for a single repeat).

I14: Spacer hits were detected to multiple repeats assigned to Bacillota_A, *Bacillota*, and *Pseudomonadota* with high confidence. Spacer hits were also detected to two repeats assigned to Patescibacteria, but this assignment was made with low confidence (repeats assigned based on detection in a single MAG). A relatively low number of hits were obtained for repeats assigned to *Pseudomonadota* or *Bacillota* (≤9, and ≤5 with 0 or 1 mismatches), while a larger number of hits were obtained for repeats assigned to Bacillota_A (up to 47, up to 14 hits with 0 or 1 mismatches).

I49: Spacer hits were obtained from 3 repeats, all with a low-confidence taxonomic assignment (taxonomic assignment based on detection of the repeat in a single MAG).

I60: Spacer hits were detected from multiple repeats assigned to Desulfobacterota, including one with high confidence. The number of hits for these repeats was also relatively high (multiple repeats with ≥80 hits with 0 or 1 mismatch). Spacer hits were also detected for a single repeat assigned to *Streptococcus* with medium confidence (repeat detected in a single isolate genome), and one assigned to *Elusimicrobiota* with low confidence. The number of hits associated with these latter two repeats was lower (<25).

I15: Spacer hits were detected from repeats assigned to *Bacillota* and Bacillota_C, one repeat with high-confidence assignment for each phylum (*Bacilli* ; *Streptococcus* and *Negativicutes* ; *Megasphaera*), as well as one repeat assigned with medium confidence to *Halobacteriota* (*Methanomicrobia* ; *Methanofollis*). Most individual repeats showed a relatively low number of spacer hits, with ≤2 hits with 0 or 1 mismatch for the two repeats with high-confidence assignment.

I86: Spacer hits were detected to one repeat assigned to *Halobacteriota* with medium confidence, as well as single repeats assigned to *Pseudomonadota* or *Verrucomicrobiota* with low confidence. All repeats showed a relatively high number of hits (>50 hits), the vast majority without any mismatch between spacer and target.

I87: Spacer hits were detected from multiple repeats assigned to *Methanoculleus* with medium confidence. The number of this for these repeats was relatively high (multiple repeats with ≥50 hits with 0 or 1 mismatch). Spacer hits were also detected for a single repeat assigned to *Thermoplasmatota* and a single repeat assigned to *Thermotogota*, both with low confidence. The number of hits associated with the *Thermotogota* repeat was also high (>100 hits) while this number was lower for the *Thermoplasmatota* repeat (8 hits).

I10: Spacer hits were detected from multiple repeats assigned to *Halobacteriota* with medium confidence, and for one repeat assigned to Bacillota_A with low confidence (repeat detected in a single MAG). A large (≥500) number of hits with 0 or 1 mismatch were detected for repeats assigned to *Halobacteriota*, and a lower but still substantial number (45) were detected for the repeat assigned to Bacillota_A.

I48: All spacer hits were detected for repeats with low-confidence taxonomic assignment (repeats detected in a single MAG). A large (≥30) number of hits with 0 or 1 mismatches were detected for several repeats.

I53: Spacer hits were detected to multiple repeats assigned to *Bacteroidota* with medium confidence, as well as a single repeat assigned to *Armatimonadota* with low confidence. Both repeats assigned to *Bacteroidota* and *Armatimonadota* showed a relatively high number of hits (≥30).

**Supplementary Tables (legends)**

Table S1. **PAM motifs predictions for individual repeats based on conserved neighborhood of hits to IMG/VR and IMG/PR sequences.** Only hits with 0 or 1 mismatch between spacer and predicted protospacer were considered. Only repeats with ≥50 distinct neighborhoods (10 bases upstream and downstream of the hit) and repeats with <50 distinct neighborhoods but a predicted PAM corresponding to a known motif for their predicted type are included in this supplementary table. All motifs are shown as if upstream from the protospacer, as the spacer orientation can be ambiguous in our dataset. This means that some of these PAMs are not the exact sequence shown here, but instead the reverse-complement downstream from the spacer.

Table S2. **Functional annotation of spacer hits for different types of spacers (single vOTU vs multivOTU).** For each type of spacer (either spacers hitting a single vOTU, or spacers hitting multiple vOTUs), 8 random subsamples of 100,000 spacers were collected and functionally annotated when overlapping a CDS. The table shows the average and individual percentage of each functional category and match type (intergenic, intragenic, or overlapping gene edge) for each subsample for single vOTU spacers first, then for multivOTU spacers.

Table S3. **Targeting patterns for selected UViGs connected to different phyla.** The connection between UViGs and arrays depicted as virus-taxon networks in Fig. 6 c and d are detailed here for each individual repeat. The confidence of the taxonomic assignment is indicated along with the percentage of target covered by spacers, the number of spacer hits at different mismatch levels (from 0 to 3) and the corresponding correlation level between spacer hit number and mismatch number.

**Supplementary Data Files (list and links)**

Data file S1. **List of repeat sequences used to mine CRISPR spacers in metagenomes.** https://portal.nersc.gov/dna/microbial/prokpubs/spacer_database_resources/Supplementary_Data_File_1_repeats.tsv

Data file S2. **List of metagenomes mined for CRISPR spacers.** https://portal.nersc.gov/dna/microbial/prokpubs/spacer_database_resources/Supplementary_Data_File_2_samples.tsv

### Supplementary Figures

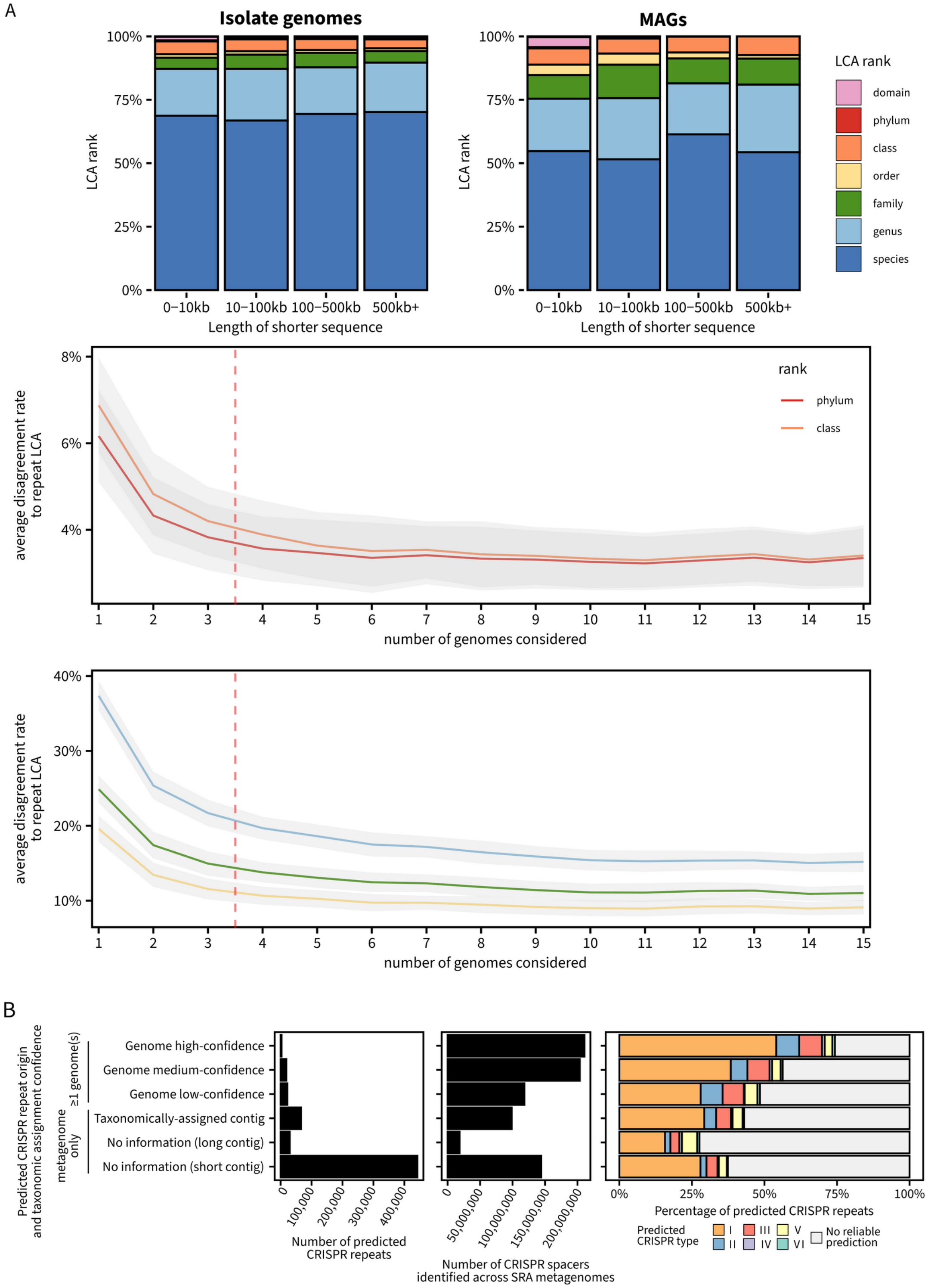

Figure S1. **Taxonomic and type distribution of predicted CRISPR repeats.** A. The top panel shows the lowest common ancestor (LCA) rank for individual CRISPR repeats detected in ≥2 genomes for isolate genomes (left) and metagenome-assembled genomes (MAGs, right). The bottom panels show the consistency in predicted LCA when considering only a subset of the genomes (x-axis) at different taxonomic ranks (colors). The orange dashed line indicates the cutoff for minimum number of genomes (≥4) required to consider a taxonomic assignment as "high-confidence" (if isolates) or "medium-confidence" (if MAGs, see Methods). B. Number of repeats and associated spacers for each category of repeat, i.e. repeats identified in genomes (with different levels of taxonomic assignment confidence), or repeats identified exclusively in metagenome contigs. The rightmost bar chart shows the distribution of predicted CRISPR types for each repeat category.

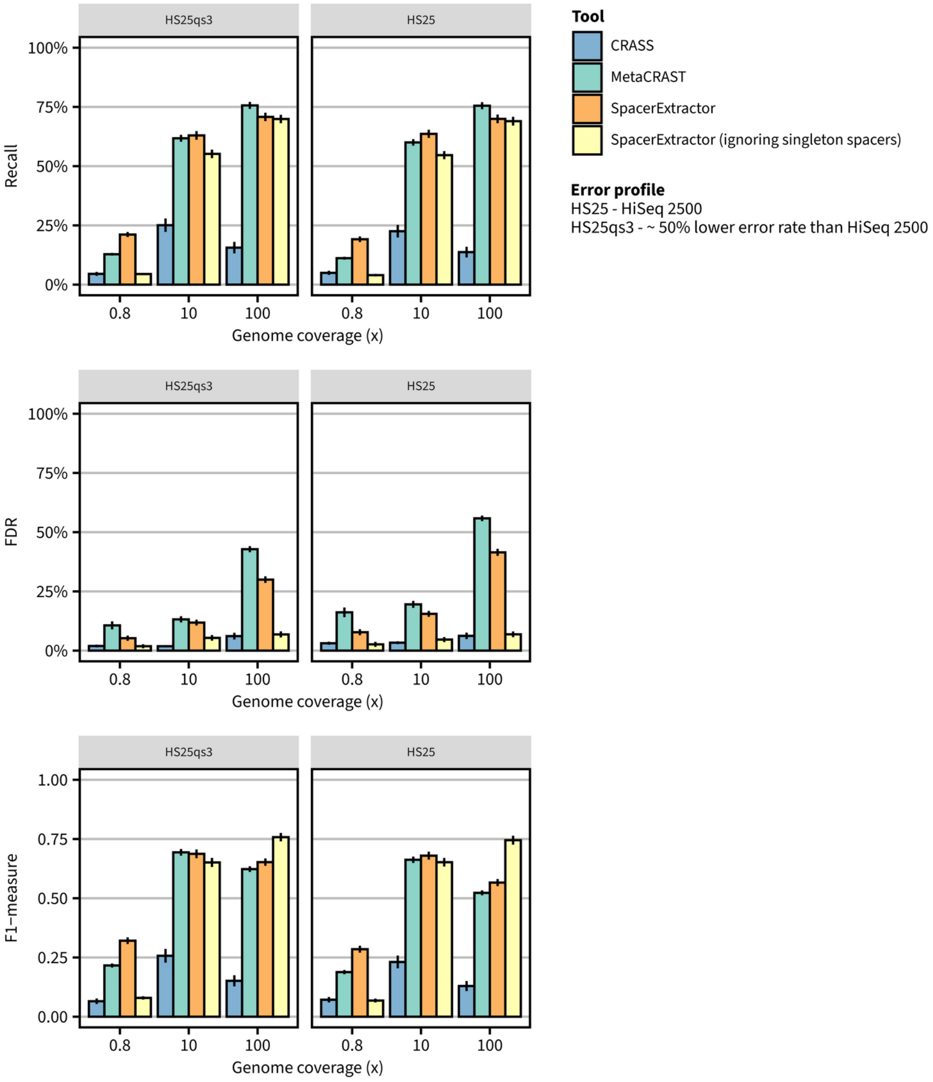

Figure S2. **Evaluation of spacer recovery from short reads across coverage, error rate, and tools**. Three metrics were computed based on the prediction of spacers by CRASS, MetaCRAST, and SpacerExtractor: recall (top row), false-discovery rate (middle row), and F1 score (bottom row). For SpacerExtractor, these metrics were calculated both with all high-quality spacers, or with all high-quality spacers excluding singletons, i.e. requiring spacer coverage >1. Two error profiles were used when simulating reads, corresponding to ~50% of the error rate of the Illumina HiSeq 2500 sequencer (left column) or ~100% of the error rate of the Illumina HiSeq 2500 sequencer (right column), and three levels of genome coverage were used (0.8x, 10x, 100x, x-axis). For each metric, the average and standard error across 200 genomes is indicated.

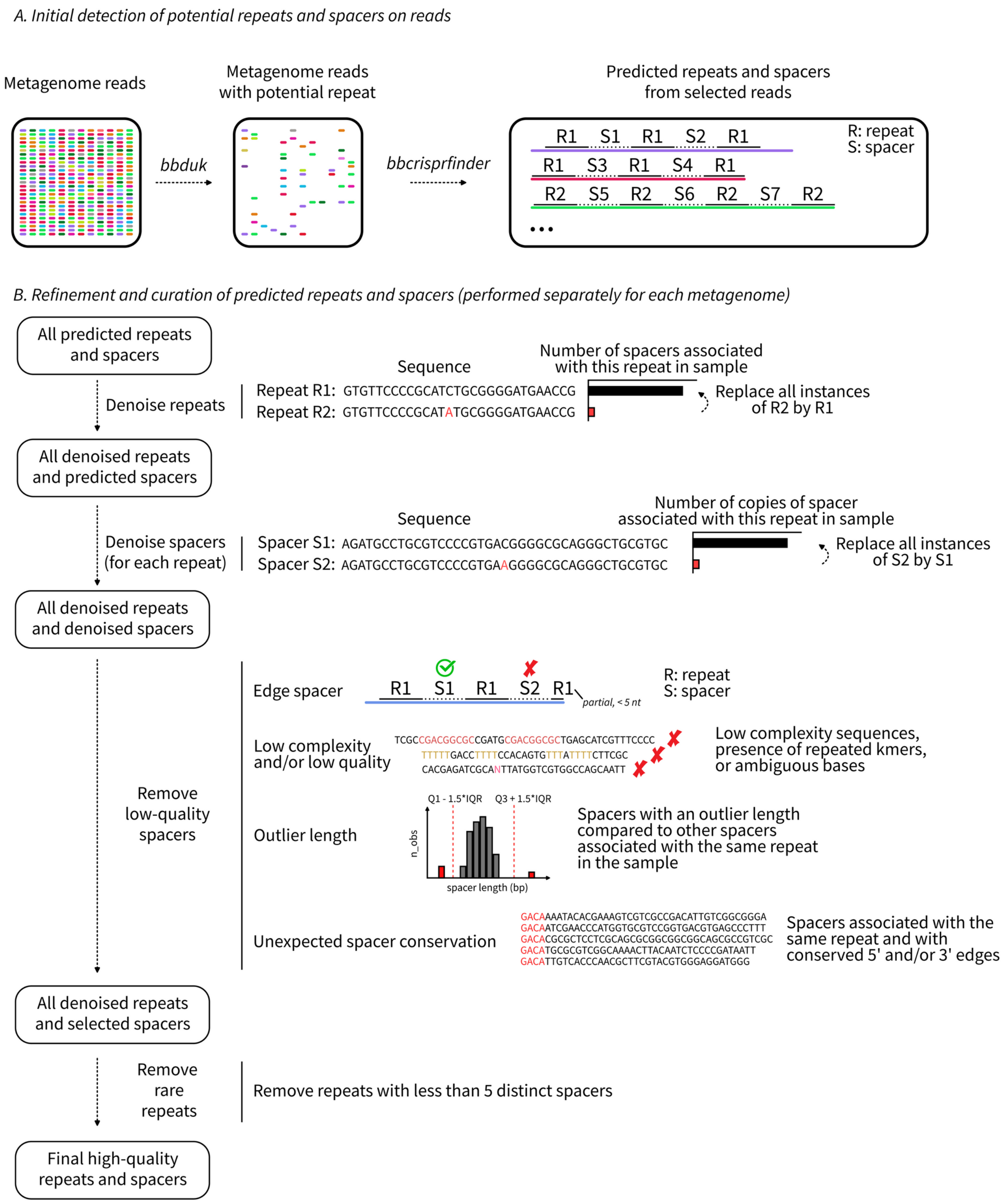

Figure S3. **Schematic overview of the pipeline used to collect spacers from metagenomic reads.** These steps correspond to the SpacerExtractor pipeline v0.8 with default parameters, including a requirement of 5 spacers without any quality warning for a given repeat to be considered as reliably detected in a metagenome, and the corresponding spacers to be considered as potentially high-quality (depending on the quality warnings).

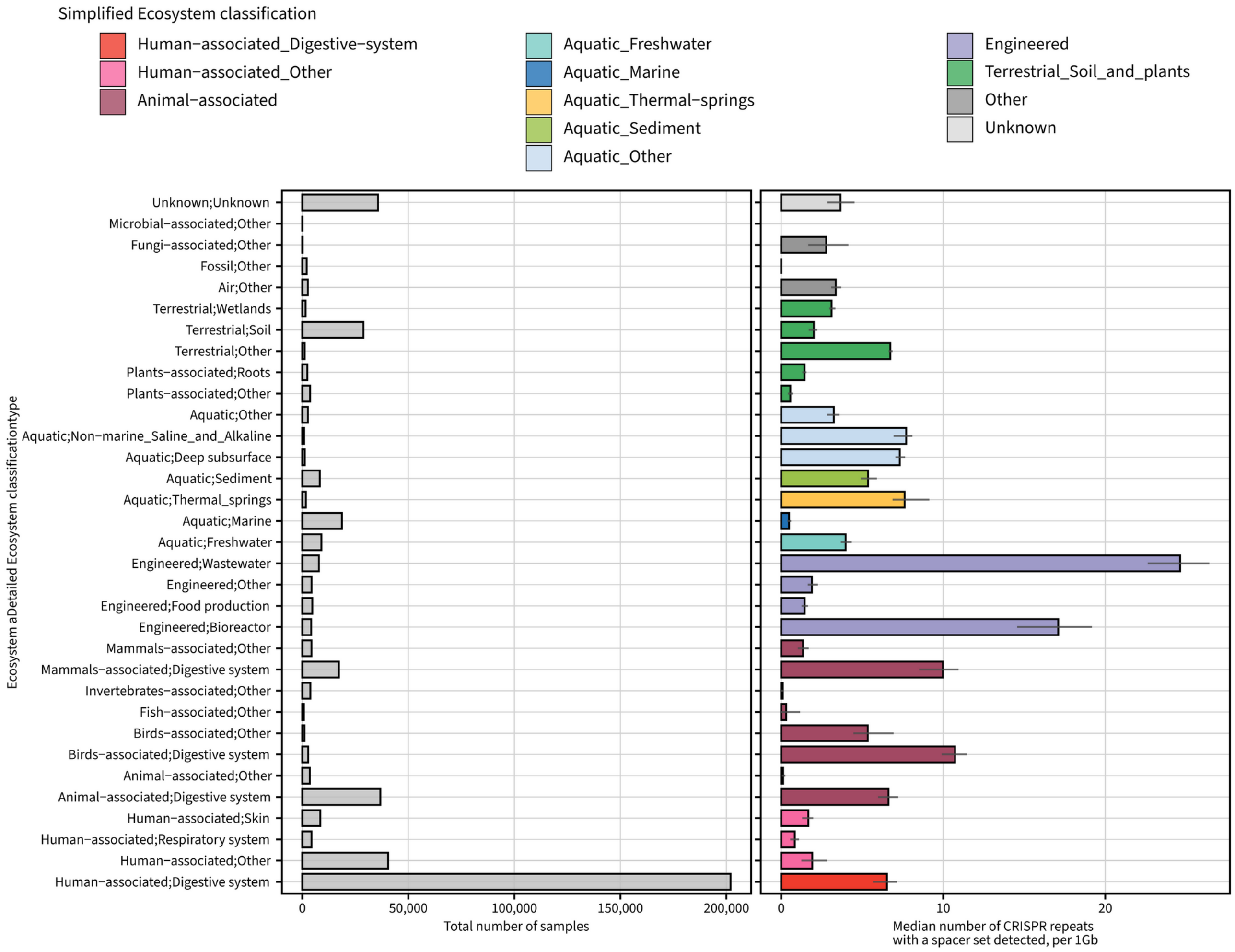

Figure S4. **Number of distinct CRISPR repeats detected per sample across ecosystems**. The left panel shows the total number of samples for each ecosystem type (based on the detailed ecosystem classification). The right panel shows the median number of CRISPR repeats with a detected spacer set, normalized per 1Gb of metagenomes. Median numbers were calculated for 50 random subsamples of 1,000 samples, 500 samples, or 50 samples, depending on the total number of samples with 0.5Gb of data or more available for the category (the largest subsample possible was selected for each category). The bar chart shows the average value obtained across the subsamples, and the error bar represents the minimum and maximum values observed across subsamples.

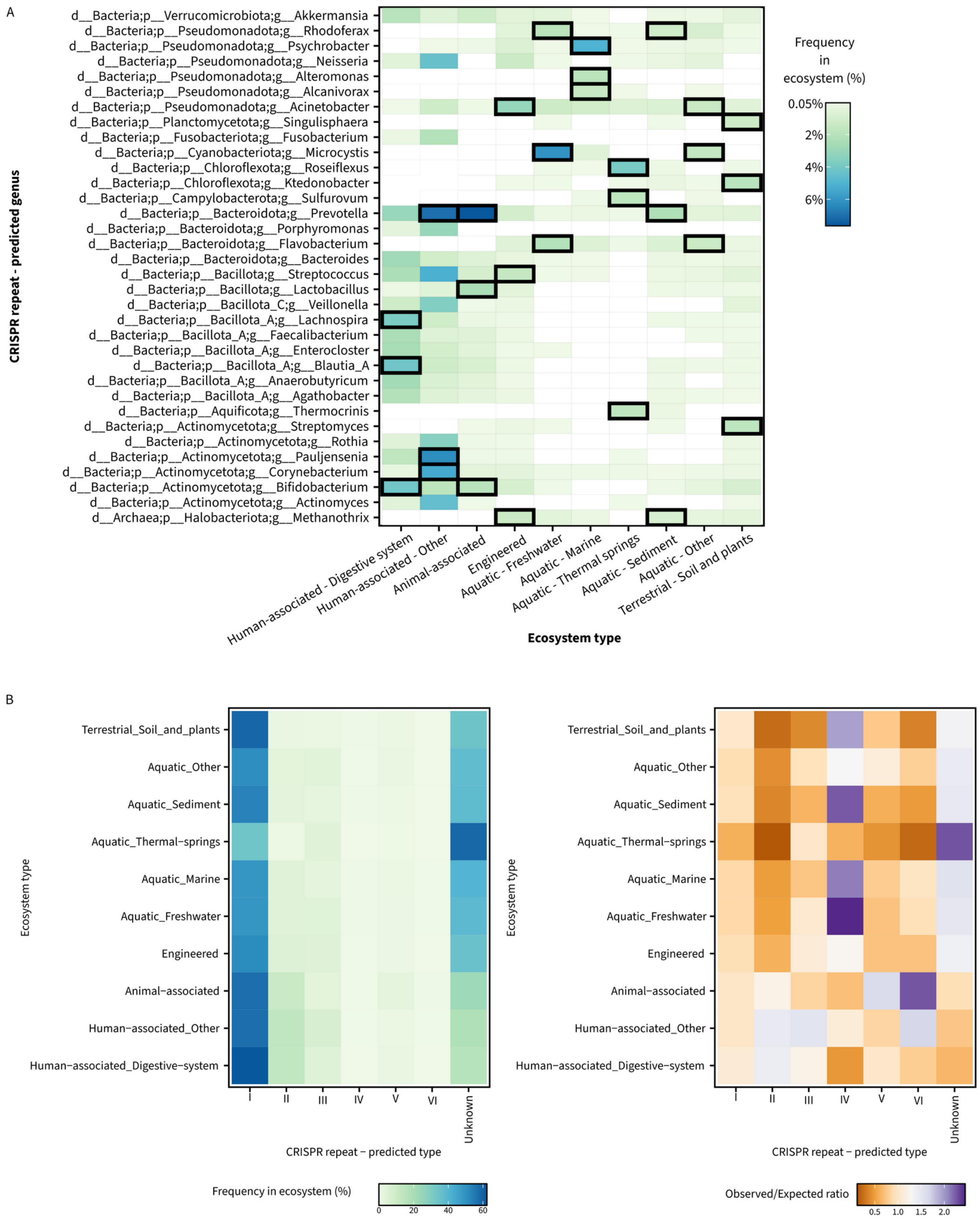

Figure S5. **Taxonomic assignment and predicted type of CRISPR repeats detected by ecosystem**. A. For each ecosystem group (x-axis), each taxon relative frequency was calculated among all CRISPR repeats with a taxonomic assignment to the genus or species rank. For the heatmap display, all taxa with a frequency ≥2% or within the top 3 taxa for an ecosystem were included, and all frequencies <0.05% were set as 0 for clarity. The 3 most frequently detected taxa for each ecosystem are highlighted with a black outline. B. The left heatmap shows the frequency of CRISPR type (columns) across all CRISPR repeats with a detected spacer set for each ecosystem (rows). “Unknown” CRISPR types are predicted CRISPR repeats for which no type could be predicted. The right panel shows the same data in the form of an “observed:expected” ratio, with the expected counts calculated based on a uniform distribution of CRISPR types across ecosystem types.

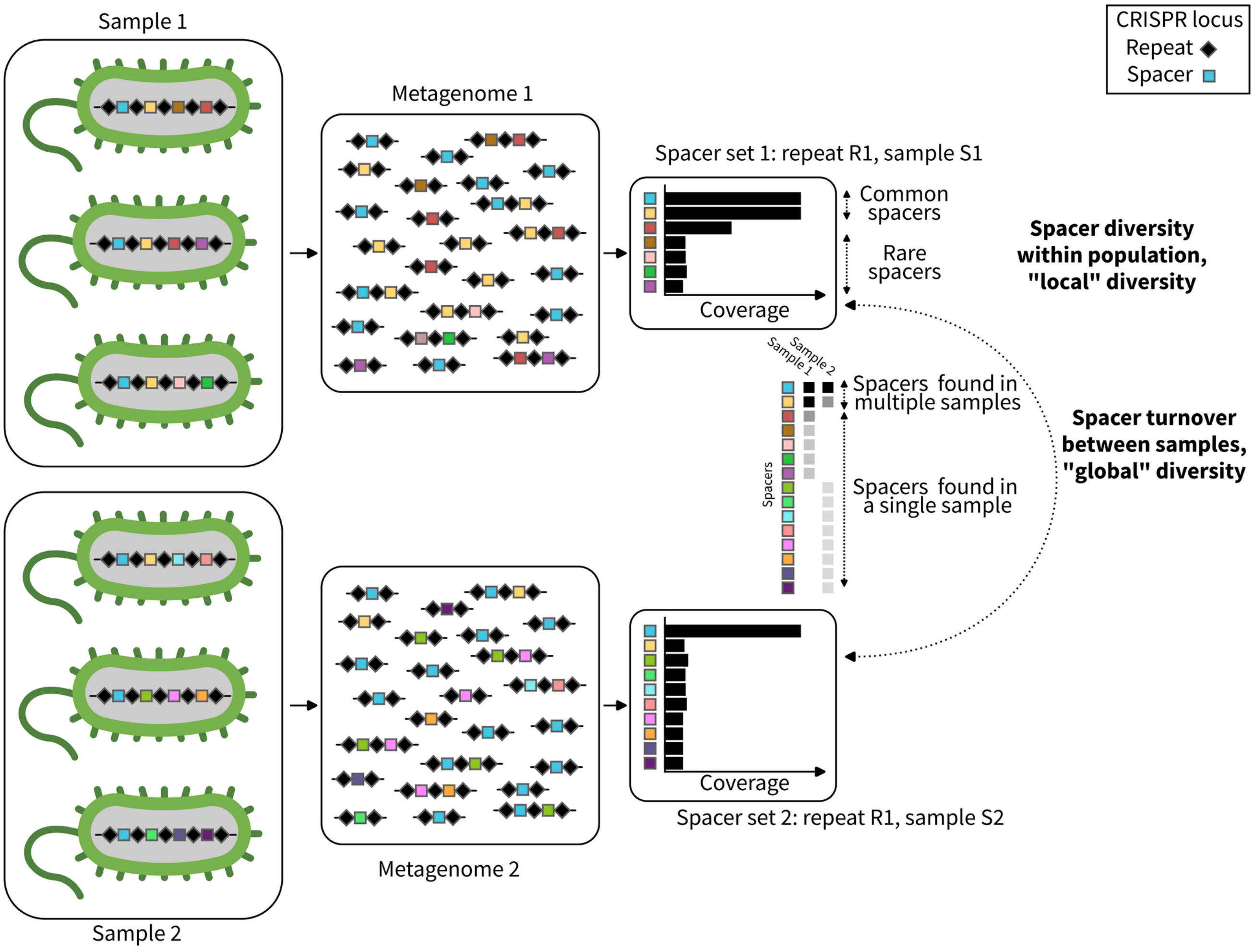

295 Figure S6. **Schematic overview of the spacer diversity analysis**. For individual repeat sequences, the coverage of spacers obtained from metagenomic reads was compared within sample to identify common and rare spacers, and between samples to evaluate spacer turnover.

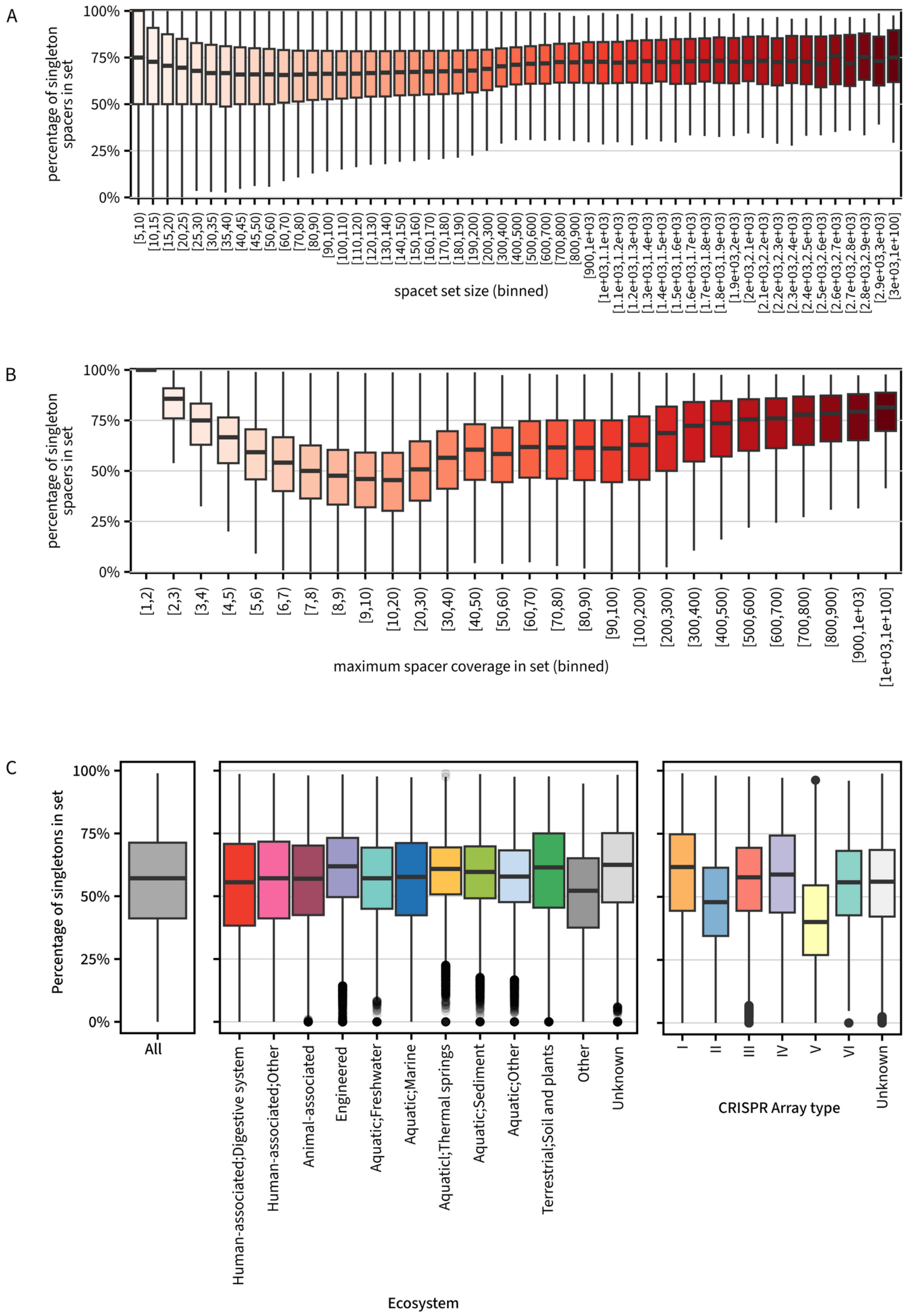

Figure S7. **Percentage of singletons across spacer set size, maximum spacer coverage in set, ecosystem, and CRISPR array type.** A. Relationship between the percentage of singletons in a spacer set (y-axis) and the size of spacer sets (x-axis). B. Relationship between the percentage of singletons in a spacer set (y-axis) and the maximum spacer coverage in the spacer set (x-axis). C. Distribution of the percentage of singletons in a spacer set (y-axis) for all sets (left), by ecosystem (middle), and CRISPR array type (right). Only spacer sets for which the maximum spacer coverage was 20x or higher were included in plots for the bottom row.

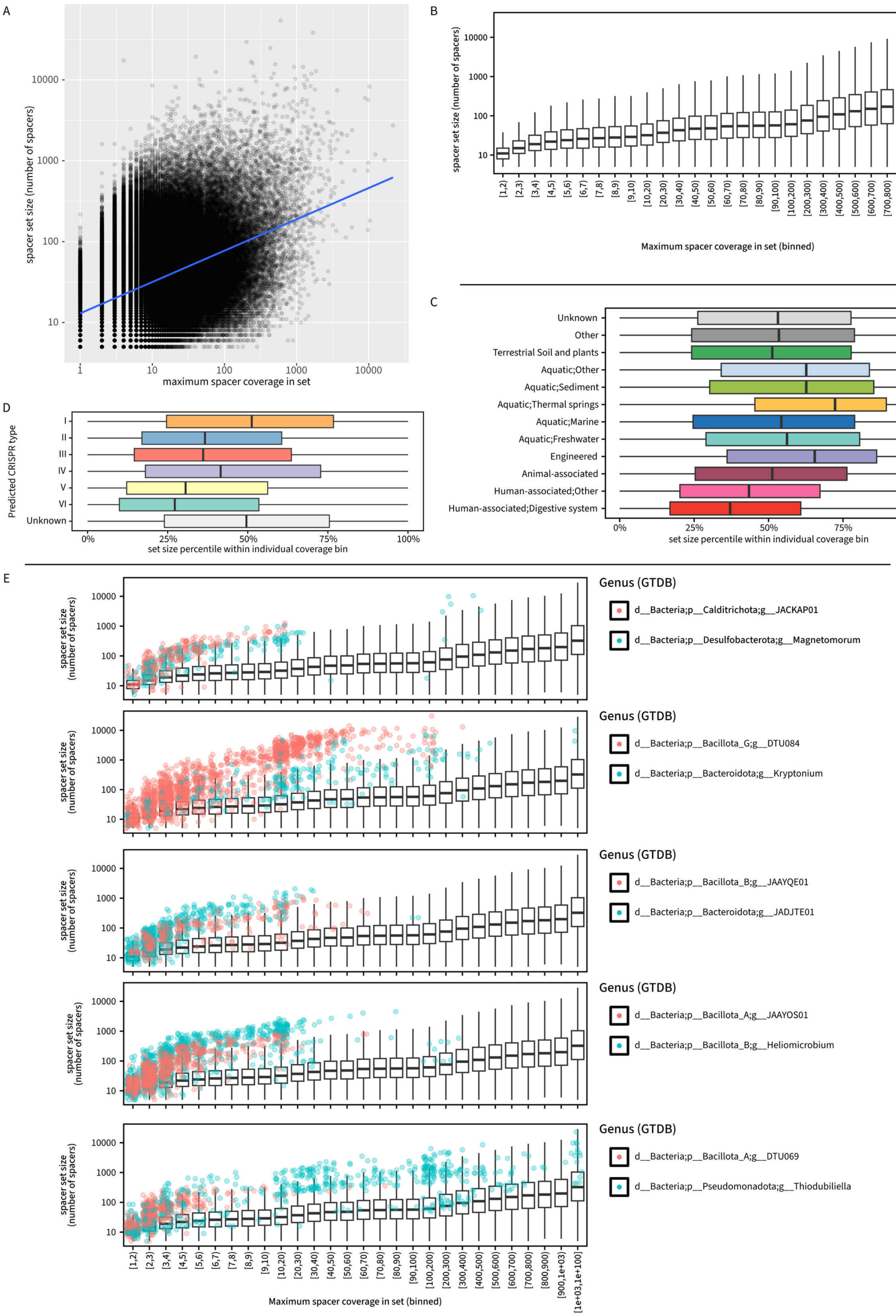
A
spacer set size (number of spacers)
maximum spacer coverage in set
B
spacer set size (number of spacers)
Maximum spacer coverage in set (binned)
C
Unknown
Other
Terrestrial Soil and plants
Aquatic;Other
Aquatic;Sediment
Aquatic;Thermal springs
Aquatic;Marine
Aquatic;Freshwater
Engineered
Animal-associated
Human-associated;Other
Human-associated;Digestive system
set size percentile within individual coverage bin
D
Predicted CRISPR type
I
II
III
IV
V
VI
Unknown
set size percentile within individual coverage bin
E
spacer set size (number of spacers)
Genus (GTDB)
d__Bacteria;p__Calditrichota;g__JACKAP01
d__Bacteria;p__Desulfobacterota;g__Magnetomorum
d__Bacteria;p__Bacillota_G;g__DTU084
d__Bacteria;p__Bacteroidota;g__Kryptonium
d__Bacteria;p__Bacillota_B;g__JAAYQE01
d__Bacteria;p__Bacteroidota;g__JADJTE01
d__Bacteria;p__Bacillota_A;g__JAAYOS01
d__Bacteria;p__Bacillota_B;g__Heliomicrobium
d__Bacteria;p__Bacillota_A;g__DTU069
d__Bacteria;p__Pseudomonadota;g__Thiodubiliella
Maximum spacer coverage in set (binned)

Figure S8. **Spacer set diversity as a function of spacer set size, ecosystem, and CRISPR array type. A.** Relationship between spacer set size (y-axis) and maximum spacer coverage in a set (x-axis), on a log10-log10 scale. The dots are a random subset of 500,000 samples, while the blue line shows the results of a linear regression over all samples. **B.** Distribution of spacer set size (y-axis) for sets grouped by maximum spacer coverage in set (x-axis). **C.** Distribution of the set size percentiles within their coverage bin across ecosystems. For each set, the set size was transformed in a percentile when compared to all other sets within the same coverage bin (see panel B). The distribution of percentiles is then plotted by ecosystem, to highlight ecosystems for which spacer sets tend to be always on the larger or smaller end of the distribution within their coverage bin. **D.** Distribution of set size percentiles for CRISPR array types. **E.** Distribution of set size for the 10 genera with the largest percentage of sets in the 80$^{th}$ size percentile or above. For each plot, the distribution of set sizes for all taxa is indicated by a boxplot, and the set size corresponding to the genera selected are indicated with colored dots (2 per plot).

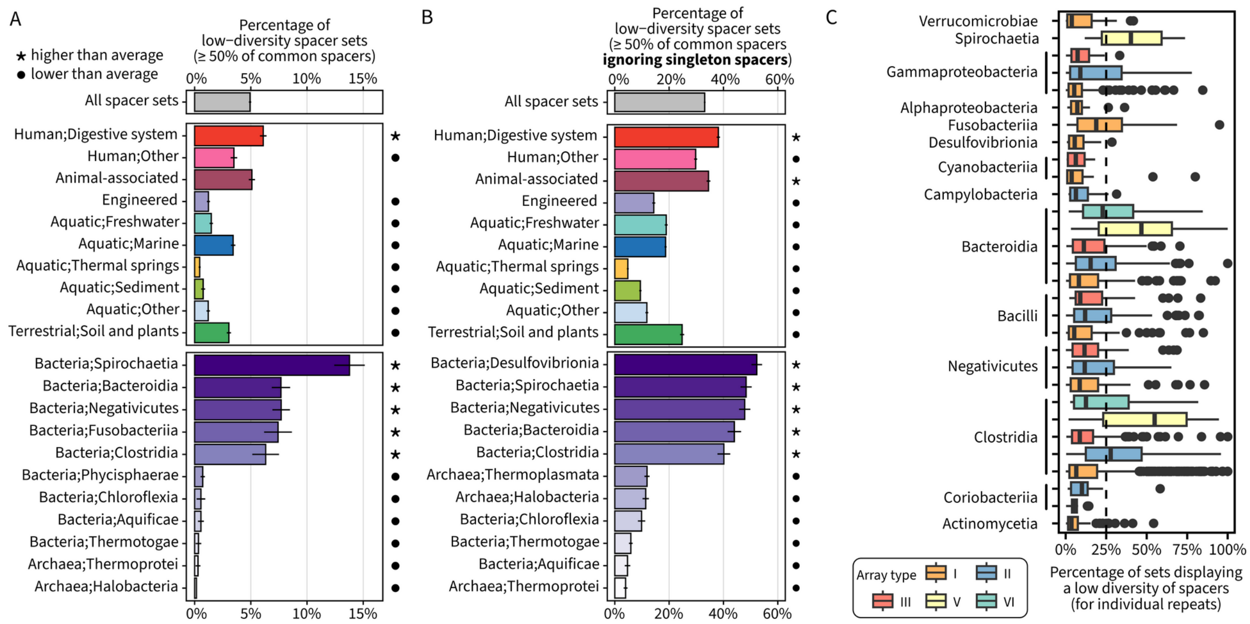

Figure S9. **Overview of low-diversity spacer sets distribution. A.** Percentage of low-diversity spacer sets for all sets (top), by ecosystem (middle), or by class (bottom). Spacer sets are considered as low-diversity if ≥50% of the spacers are considered as "common" in the population, i.e. their coverage depth is ≥50% of the maximum coverage in the set. Only spacer sets for which the maximum coverage was ≥20x were considered. For the taxa plot, only the 5 taxa with the highest and lowest percentage of low-diversity spacer sets are shown. Z-tests with a p-value cutoff of 0.001 were used to identify percentages significantly higher or lower than average (stars and circle symbols, respectively). **B.** Percentage of low-diversity spacer sets similar as panel A, but calculated without considering singleton spacers, i.e. the percentage of common spacers is calculated only among non-singleton spacers. **C.** Distribution of the percentage of samples in which a spacer set was identified as low diversity (≥50% of common spacers, x-axis), per taxon and CRISPR array type (y-axis). Only arrays with at least 1 low-diversity sample and ≥10 samples with maximum spacer coverage ≥20x were considered, and only taxon-type combinations with 10 distinct repeats were included in the plot.

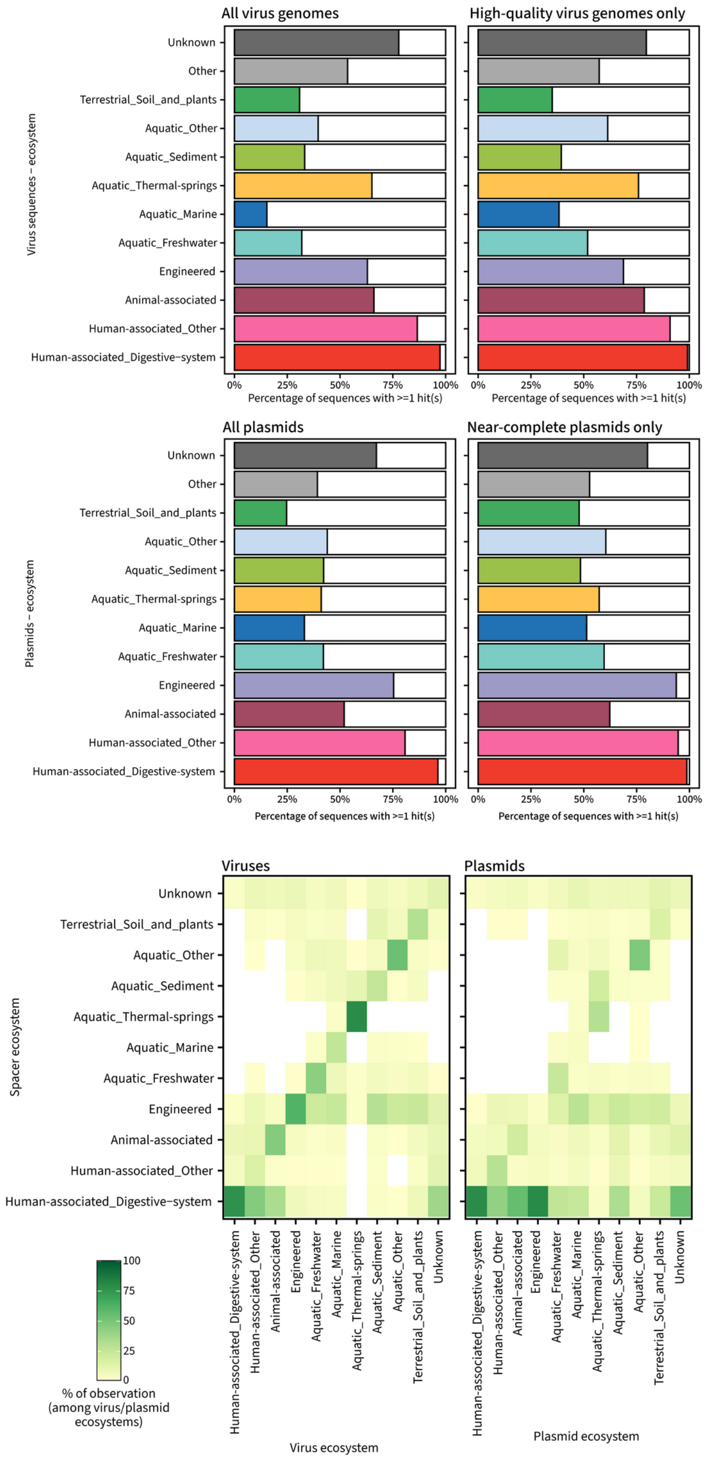

335 Figure S10. **Spacer hits to IMG/VR and IMG/PR across ecosystems.** The percentage of targets (top panel: viruses, middle panel: plasmids) with at least 1 hit in the global spacer database, based on the ecosystem from which the corresponding target was assembled from (y-axis). For each target, the left panel has all sequences (viruses or plasmids), and the right panel has high-quality sequences only (high-quality UViGs or near-complete plasmids). The bottom panel shows the frequency of matches organized by target ecosystem (x-axis) and spacer
340 ecosystem (y-axis). The heatmap is colored based on the percentage of hits within the target ecosystem.

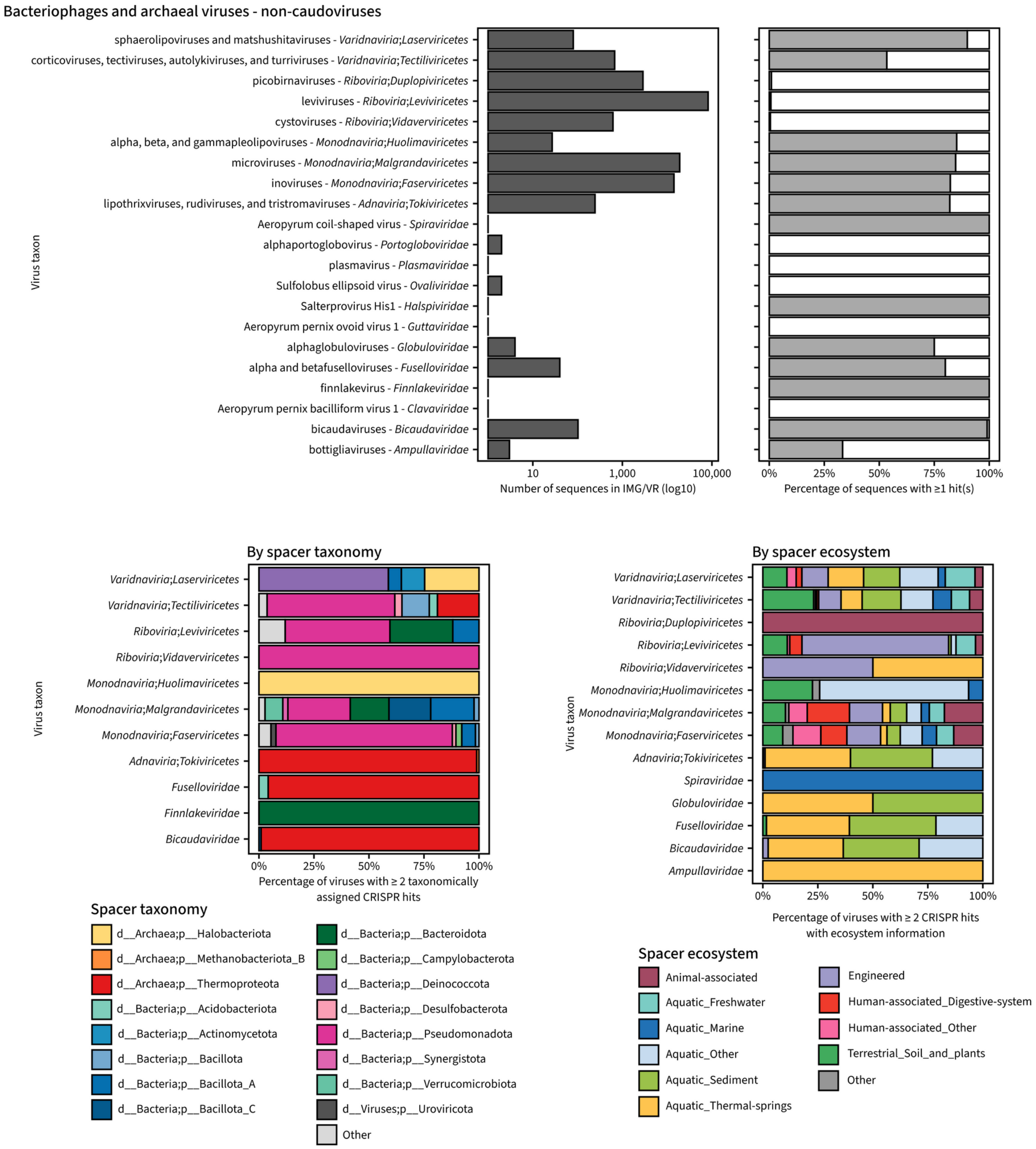

Figure S11. **Spacer hits to atypical bacteriophage and archaeal virus taxa**. The top panel shows the number of sequences in IMG/VR and the corresponding percentage of sequences with at least 1 hit in the global spacer database for taxa of bacteriophages or archaeoviruses outside of the *Caudoviricetes* class. Only taxa with a known class or family (if not classified in any known class) were included. The bottom panels show the distribution of spacer taxonomy (left) and ecosystem (right) in cases where this information is available for the corresponding hits. For the taxonomy and ecosystem distribution, a virus was connected to a taxon or ecosystem if at least 2 distinct spacer hits were observed.

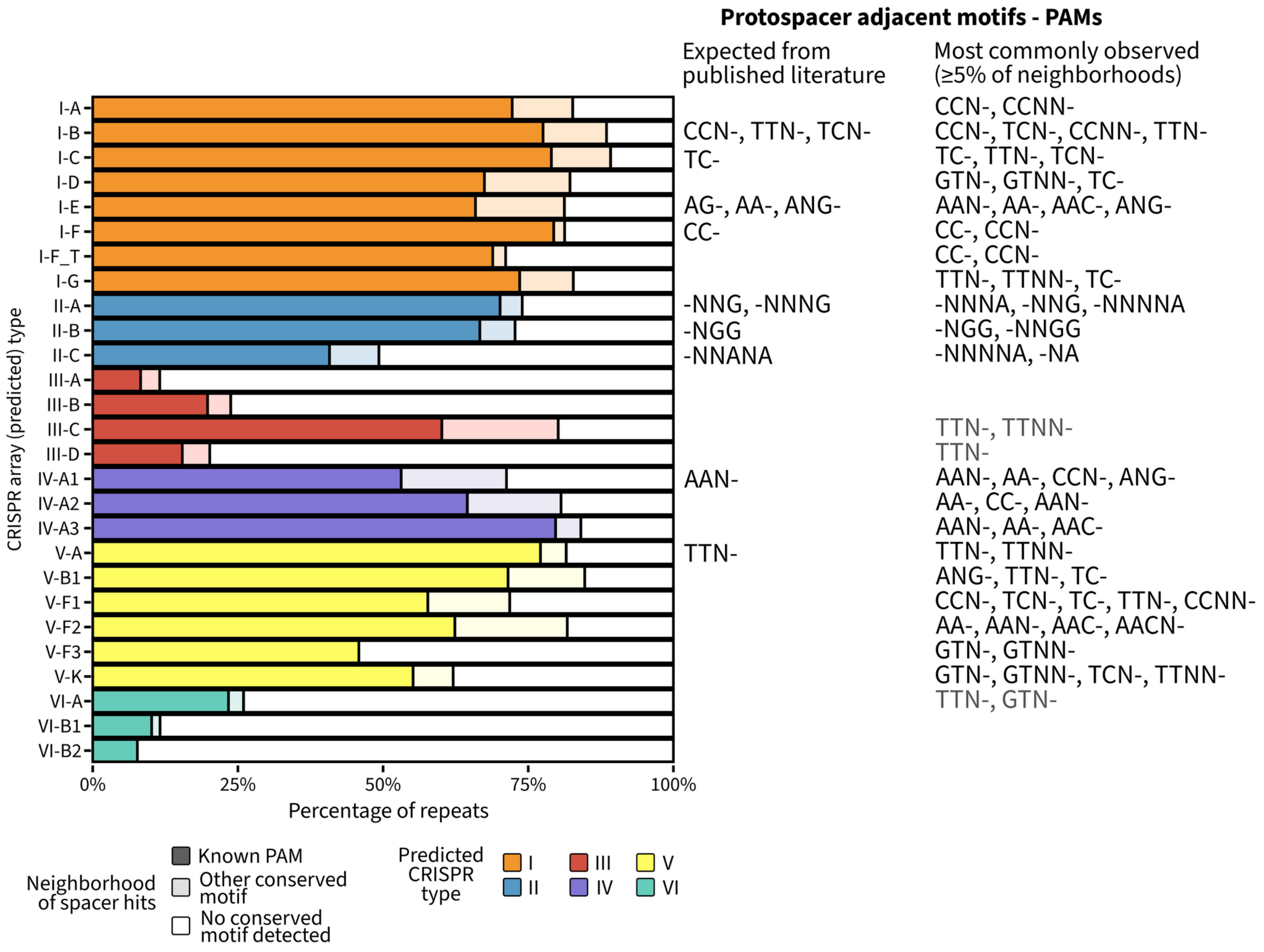

Figure S12. **Prediction of PAM motifs based on sequence conservation in spacer hits neighborhoods.** This analysis was conducted on hits detected in IMG/VR and IMG/PR sequences. Conserved positions were detected in distinct 10-bp regions upstream and downstream of spacer hits (0 or 1 mismatch), for each unique repeat. A position was considered as conserved if the same nucleotide was observed in ≥75% of the unique regions. The motifs obtained were compared to motifs previously described in the literature, and a full list of PAM motif predictions is available in Table S1.

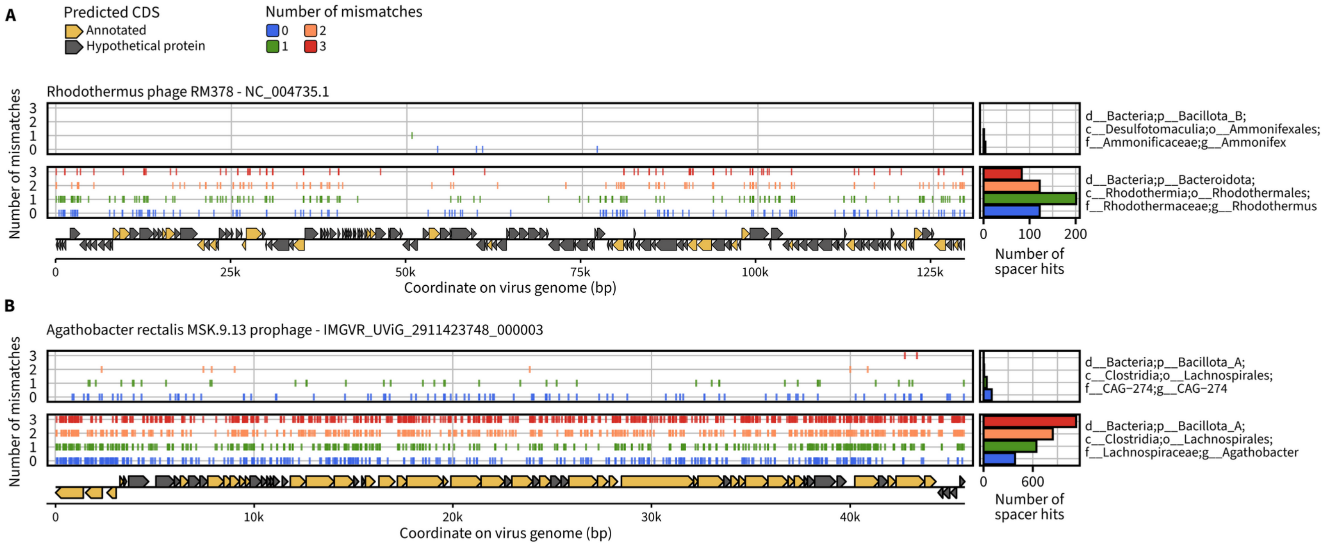

Figure S13. **Illustration of different targeting levels observed for high-quality virus genomes**. Panels A and B present an overview of spacers assigned to different taxa (indicated on the right of the plot) matching two high-quality UViGs: the genome of the isolated phage *Rhodothermus* RM378, and a prophage detected in *Agathobacter rectalis* MSK.9.13. Each time, spacer hits from a “non-host” taxon are shown in the top box, and spacer hits from the “known host” taxon are shown in the bottom box. The main plot shows the position and number of mismatches for each individual spacer, and a bar chart on the right side shows the overall distribution of the number of mismatches for the given taxon-virus pair.

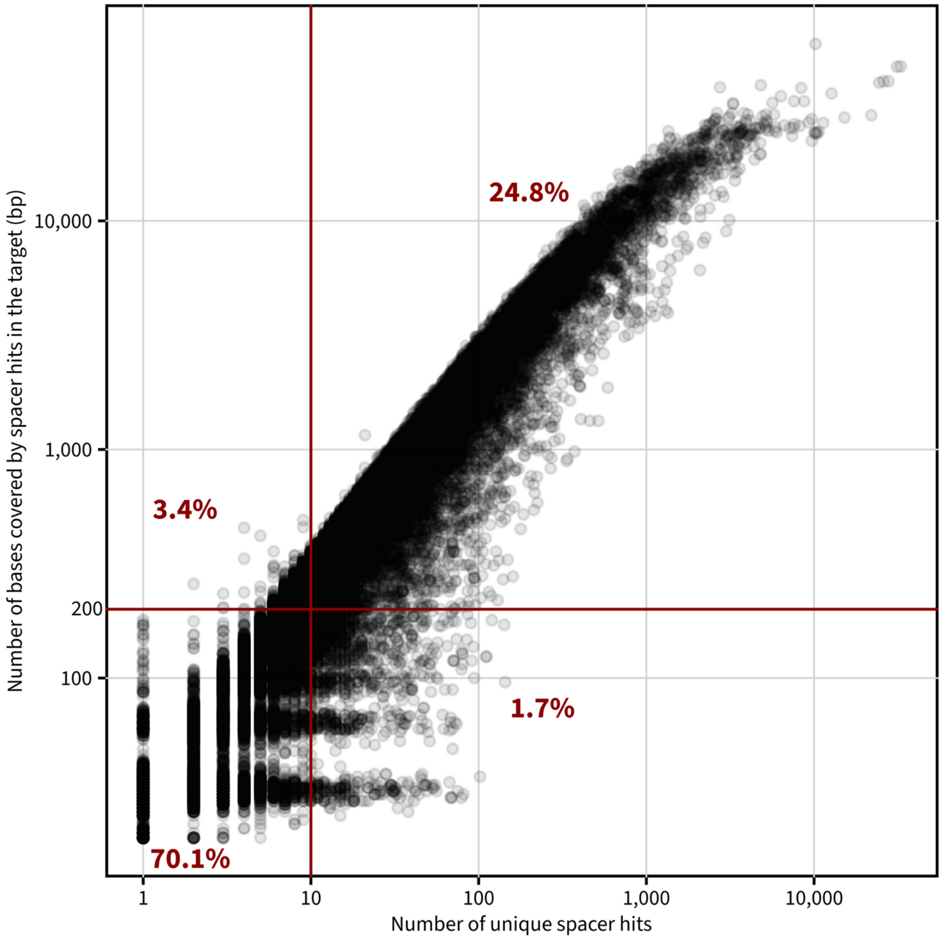

Figure S14. **Correlation between number of unique spacer hits and number of bases in the target covered by spacer hits.** The plot shows the relationship between the number of unique spacer hits (x-axis) and the number of bases covered by spacer hits (y-axis) for 100,000 randomly sampled pairs of high-quality viruses and repeat clusters. Red lines outline cutoffs of 10 distinct spacer hits and 200bp covered, and the percentage of observations within each part of the quadrant is indicated on the plot.

**I24: IMGVR_UViG_3300008155_000024**

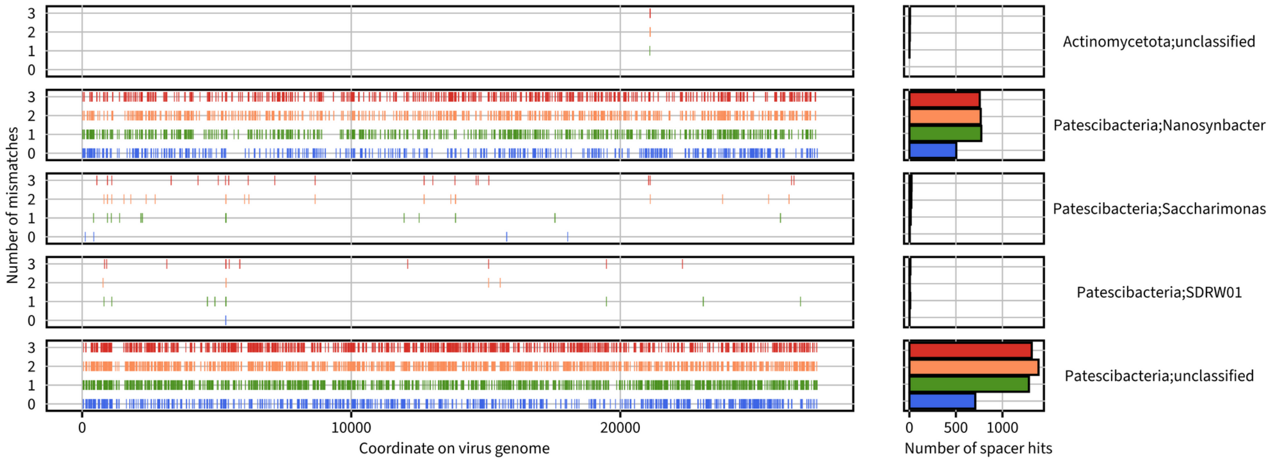

**I20: IMGVR_UViG_3300008436_000020**

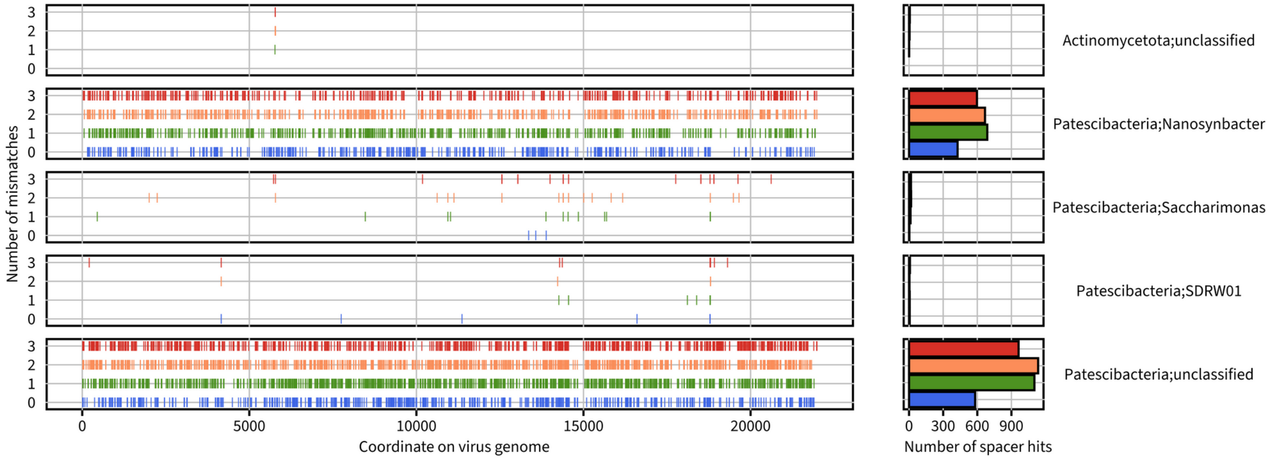

**I33: IMGVR_UViG_3300008506_000033**

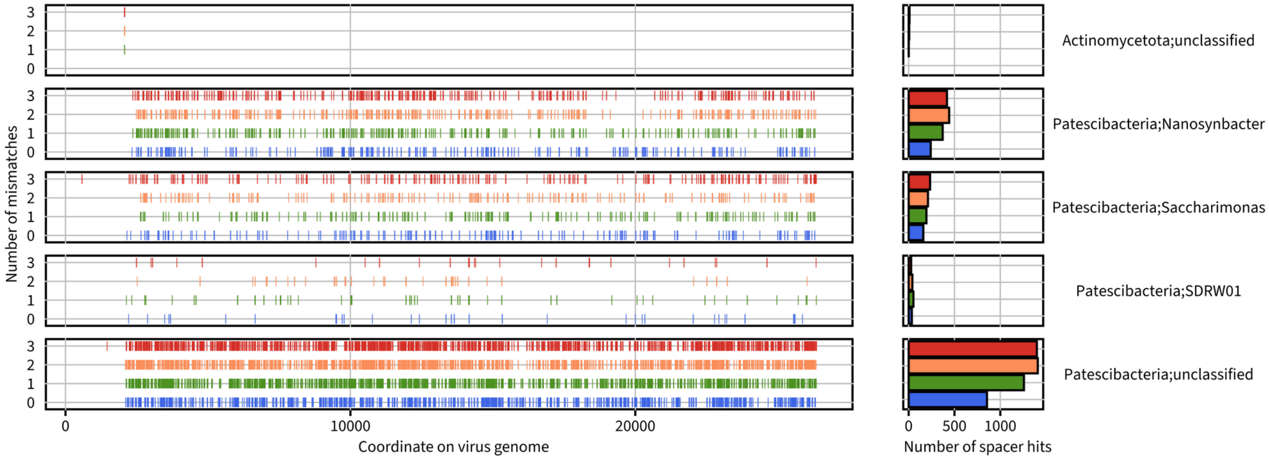

**I49: IMGVR_UViG_3300028603_000349**

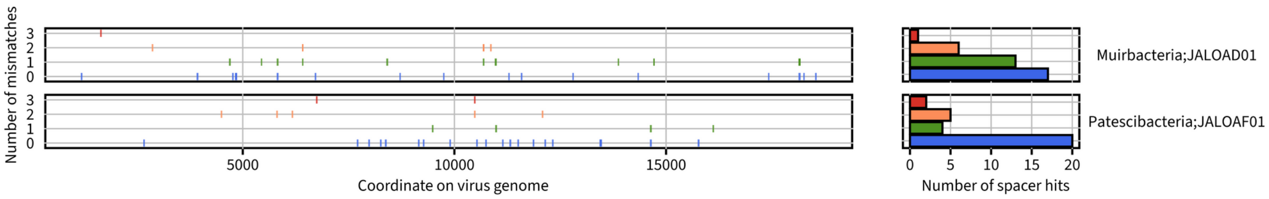

**I14: IMGVR_UViG_7000000432_000014**

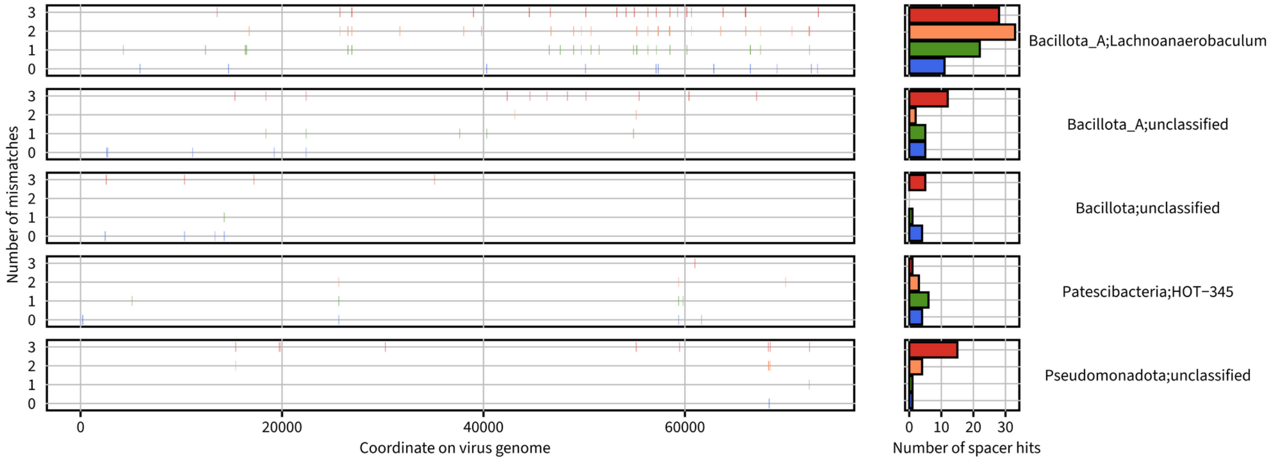

**U060: UGV−GENOME−0152060**

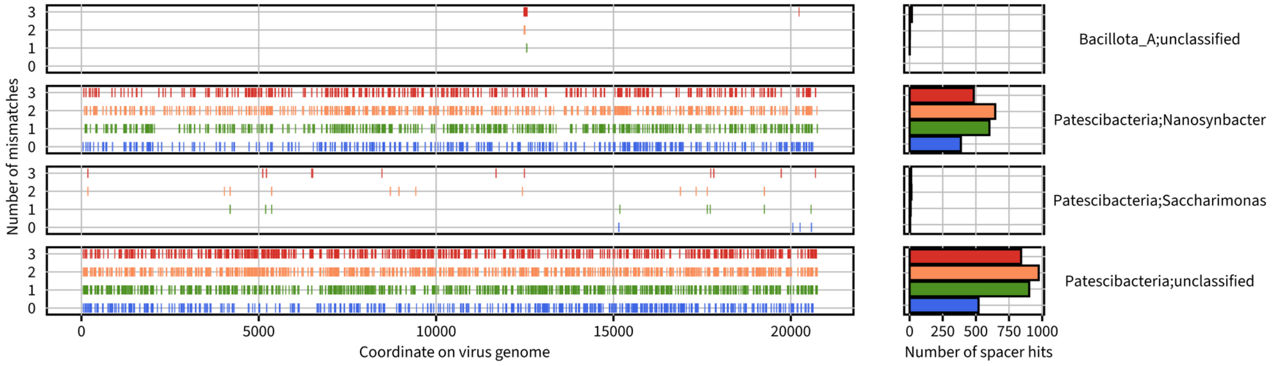

**U105: UGV−GENOME−0183105**

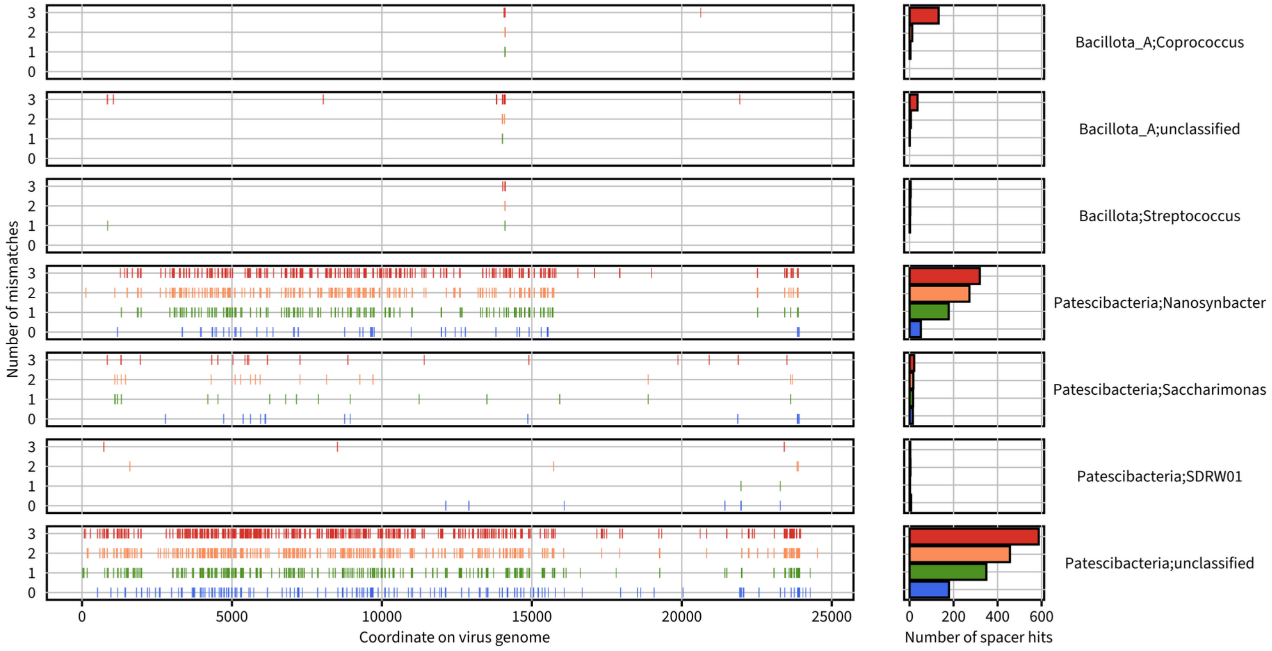

Figure S15. **Distribution of spacer hits on selected virus sequences**. For each virus sequence, the distribution of spacer hits along the virus genome (x-axis) is indicated on the left side, with the number of mismatches between spacer and virus sequences indicated on the y-axis. The right side shows the total number of spacer hits for each number of mismatches across the entire sequence. Spacer hits are gathered by predicted taxon of the corresponding repeat, at the genus rank (indicated on the right). For virus sequences, the simplified identifier is indicated next to the full identifier in IMG/VR or UGV.

**I15: IMGVR_UViG_3300014203_000115***

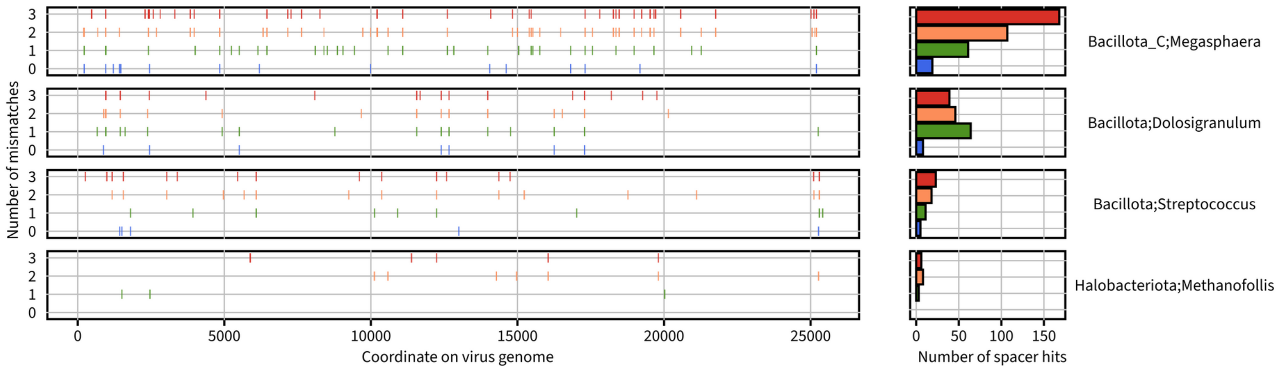

**I48: IMGVR_UViG_3300014206_000048**

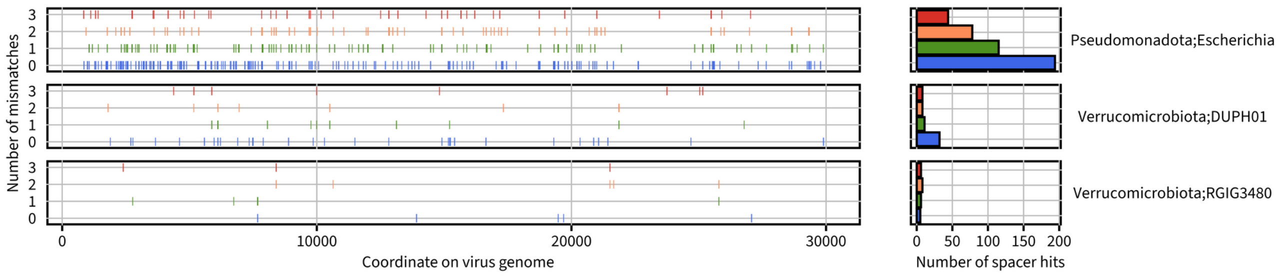

**I10: IMGVR_UViG_3300028601_000010**

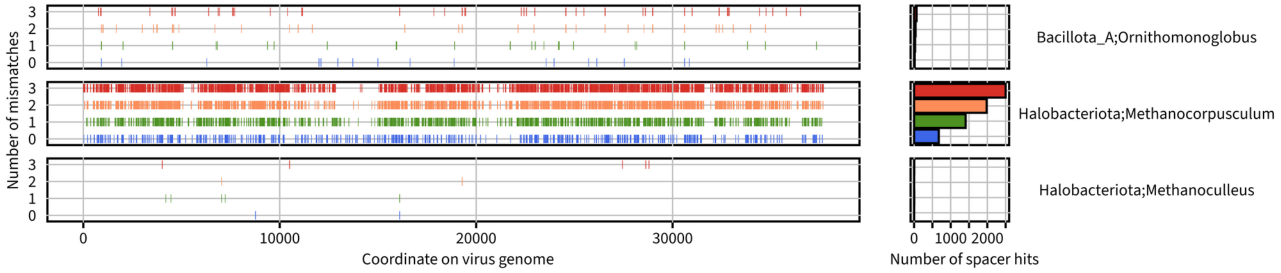

**I60: IMGVR_UViG_3300028601_000060**

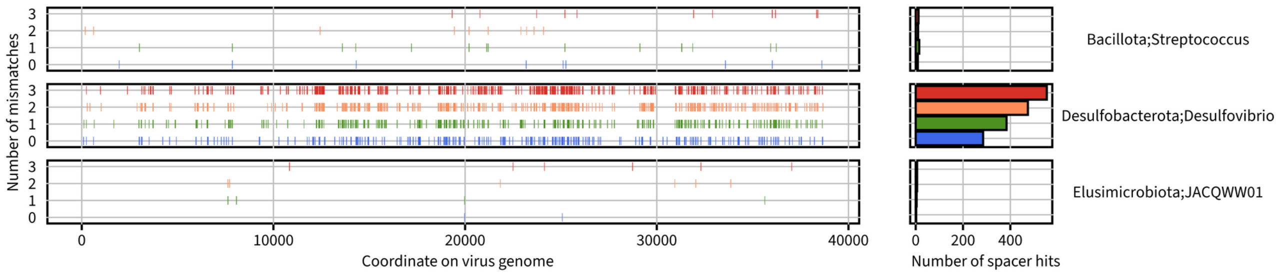

**I87: IMGVR_UViG_3300028602_002887**

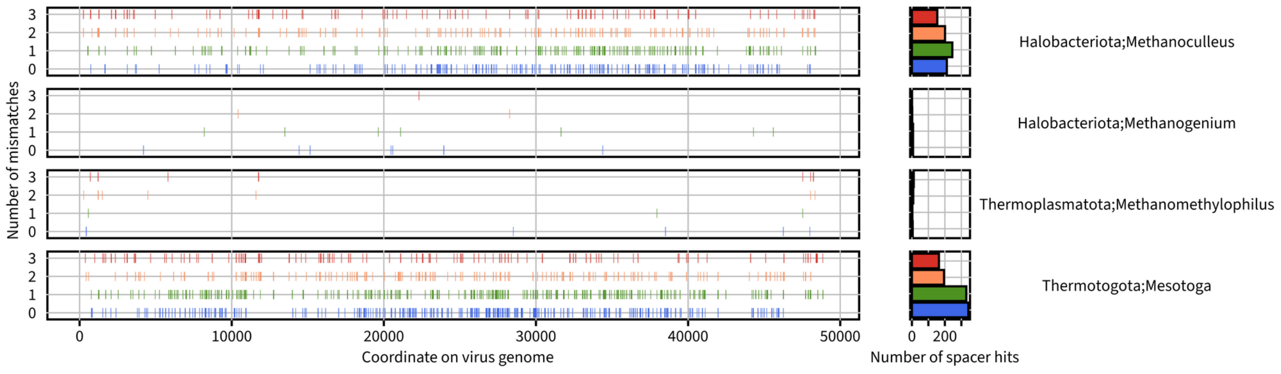

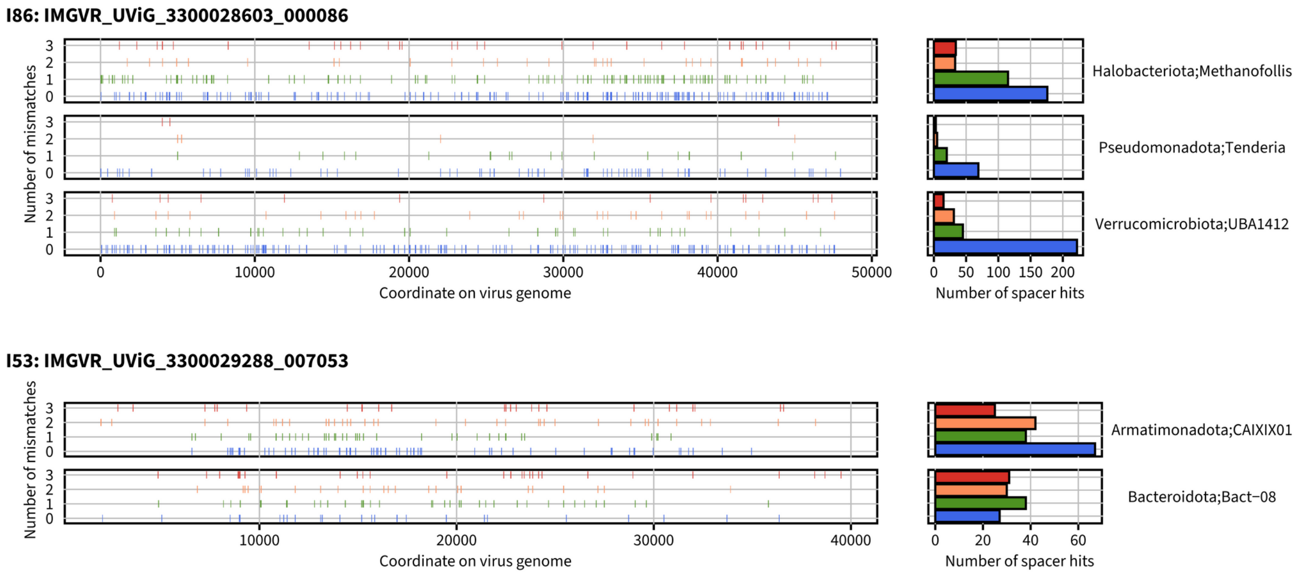

Figure S16. **Distribution of spacer hits on selected virus sequences**. For each virus sequence, the distribution of spacer hits along the virus genome (x-axis) is indicated on the left side, with the number of mismatches between spacer and virus sequences indicated on the y-axis. The right side shows the total number of spacer hits for each number of mismatches across the entire sequence. Spacer hits are gathered by predicted taxon of the corresponding repeat, at the genus rank (indicated on the right). For virus sequences, the simplified identifier is indicated next to the full identifier in IMG/VR or UGV, and an asterisk is added next to the identifier for viruses predicted to encode a DGR.

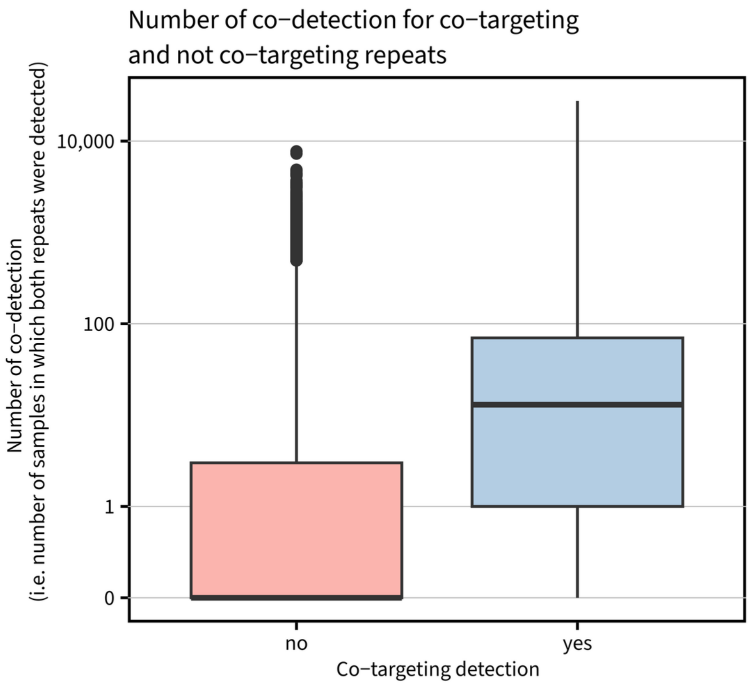

Figure S17. **Co-detection of CRISPR repeats assigned to distinct classes and targeting the same virus**. For all viruses targeted by repeats from distinct classes with ≥10 spacers, the corresponding individual repeats and their distribution across samples were collected. Next, repeat pairs were identified as co-targeting “yes” or “no” depending on whether they had at least one targeted virus in common (x-axis), and the number of samples in which both repeats were identified was tallied (y-axis). For the plot, only repeats identified in at least 100 samples were considered, and a random subsample of 350,000 repeat pairs was selected to balance the dataset between the co-targeting “yes” and “no”.

**References**

1. Skennerton, C. T., Imelfort, M. & Tyson, G. W. Crass: identification and reconstruction of CRISPR from unassembled metagenomic data. *Nucleic acids research* **41**, e105 (2013).
2. Moller, A. G. & Liang, C. MetaCRAST: reference-guided extraction of CRISPR spacers from unassembled metagenomes. *PeerJ* **5**, e3788 (2017).
3. Russel, J., Pinilla-Redondo, R., Mayo-Muñoz, D., Shah, S. A. & Sørensen, S. J. CRISPRCasTyper: Automated Identification, Annotation, and Classification of CRISPR-Cas Loci. *The CRISPR Journal* **3**, 462–469 (2020).
4. Zhang, A.-N. *et al.* CRISPR-Cas spacer acquisition is a rare event in human gut microbiome. *Cell Genomics* **0**, (2024).
5. Silas, S. *et al.* On the origin of reverse transcriptase- using CRISPR-Cas systems and their hyperdiverse, enigmatic spacer repertoires. *mBio* **8**, (2017).
6. Breusing, C. *et al.* Ecological differences among hydrothermal vent symbioses may drive contrasting patterns of symbiont population differentiation. *mSystems* **8**, e00284-23 (2023).
7. Eloe-Fadrosh, E. A. *et al.* Global metagenomic survey reveals a new bacterial candidate phylum in geothermal springs. *Nat Commun* **7**, 10476 (2016).
8. Schaible, G. A. *et al.* Multicellular magnetotactic bacteria are genetically heterogeneous consortia with metabolically differentiated cells. *PLOS Biology* **22**, e3002638 (2024).
9. Mick, E., Stern, A. & Sorek, R. Holding a grudge: Persisting anti-phage CRISPR immunity in multiple human gut microbiomes. *RNA Biology* **10**, 900–906 (2013).
10. López-Beltrán, A., Botelho, J. & Iranzo, J. Dynamics of CRISPR-mediated virus–host interactions in the human gut microbiome. *ISME J* **18**, wrae134 (2024).
11. Kirchberger, P. C. & Ochman, H. Microviruses: A World Beyond phiX174. *Annual Review of Virology* **10**, 99–118 (2023).
12. Roux, S. *et al.* Cryptic inoviruses revealed as pervasive in bacteria and archaea across Earth's biomes. *Nat Microbiol* **4**, 1895–1906 (2019).
13. Neri, U. *et al.* Expansion of the global RNA virome reveals diverse clades of bacteriophages. *Cell* **185**, 4023-4037.e18 (2022).
